# The architecture of phenotypic flexibility within a complex trait: an empirical case study using avian thermogenic performance

**DOI:** 10.1101/2021.09.27.462051

**Authors:** Maria Stager, Zachary A. Cheviron

## Abstract

Reversible modifications to trait values can allow individuals to match their phenotypes to changing environmental conditions, a phenomenon known as phenotypic flexibility. A system’s capacity for flexibility may be determined by its underlying architecture, and these relationships can have important implications for both organismal adaptation and the evolvability of acclimatization responses. Theory provides two possible alternatives to explain the ways in which lower-level traits respond to environmental challenges and contribute to phenotypic flexibility in complex, whole-organism traits: symmorphosis predicts correspondence between structure and demand across all levels of a physiological system, while the alternative predicts that influence is concentrated in select elements of a physiological network. Here we provide a rich dataset — composed of 20 sub-organismal, physiological traits paired with whole-organism metabolic rates for 106 adult Dark-eyed Juncos (*Junco hyemalis*) — to explore the mechanistic basis of phenotypic flexibility in complex traits. When exposed to synthetic temperature cues, these individuals have previously been shown to increase their thermogenic capacity (M_sum_) and enhance their ability to maintain their body temperature in the cold. We show that the relationships among a number of the traits that contribute to M_sum_ varied as the environmental context changed. Moreover, variation in M_sum_ in response to temperature acclimation was correlated with only a handful of subordinate phenotypes. As a result, avian thermogenic flexibility does not appear to be a symmorphotic response. If this is generally true of complex traits, it suggests that simple and reversible modifications can significantly impact whole-organism performance, and thus that the evolution of phenotypic flexibility in a single component part could impart flexibility for the entire system.

## INTRODUCTION

The ability to match an organism’s phenotype to changing conditions across its life can be key to fitness in variable environments (Piersma and van Gils, 2011). Such reversible modification of an individual’s trait value (phenotypic flexibility) is ubiquitous across life forms and among traits (Piersma and Drent, 2003). However, the proper matching of trait value to the demands of the environment is not guaranteed (Mills et al., 2013). Identifying the causes of variation in flexibility among individuals can therefore inform our understanding of species’ resilience to environmental change (Norin and Metcalfe, 2019). In particular, many flexible phenotypes are complex whole-organism responses that are underlain by many lower-level, subordinate traits (Schulte et al., 2011). Determining how the underlying architecture influences the system’s capacity for flexibility has important implications for understanding both organismal adaptation and the evolvability of the physiological response. For instance, in order to modify these whole-organism responses, must an individual change all subordinate phenotypes in concert or is control instead focused in just a few of these traits?

Support for concerted change derives from the evolutionary principle of symmorphosis, which states that within biological systems structural design should meet functional demand (Taylor and Weibel, 1981). This congruence between structure and function implies optimization across all levels of a physiological pathway such that no one part is operating in excess. As a result, symmorphosis predicts that parameters will exhibit an invariant ratio (i.e., constant correlations among traits) under all perturbations to the system, and empirical tests using aerobic performance have shown varying degrees of support across and within individuals (e.g., Weibel et al., 1991). However, because each component of the physiological network would need to be fine-tuned simultaneously (Dudley and Gans, 1991), this configuration could constrain the scope or rate of the flexible response.

Alternatively, we might expect that particular elements of the physiological response might be more flexible than others. In contrast to symmorphosis, this would imply that excess capacity exists in physiological systems (Diamond and Hammond, 1992). Because there are costs associated with trait modification, traits with the greatest net fitness gain should be the most flexible (Murren et al., 2015). The cost of adjusting a phenotypic value results from not only the energy directly required for trait production, but also the pleiotropic nature of many physiological traits. As with genetic pleiotropy, physiological pleiotropy can either facilitate or constrain phenotypic responses to selection (Dantzer and Swanson, 2017). It therefore follows that changing a highly pleiotropic trait may be either (1) more costly, if many downstream traits have to be changed reactively, or (2) more efficient than fine-tuning each trait individually. Depending on the structure of the physiological network, the former scenario may look much like symmorphosis. However, in the case of the latter, selection may only act on a single element to positively influence the capacity of the entire network.

Our ability to effectively evaluate these potential avenues of flexible architecture is limited by our knowledge of how organisms coordinate flexible responses in the wild. Because physiological systems are complex, it is challenging to measure all traits at once and, at the same time, traits may be responding to different environmental cues (Westneat et al., 2019). One well-studied system that lends itself to mechanistic evaluation is thermogenic flexibility — the ability to reversibly alter endogenous heat production, which is used by many small temperate birds to maintain a relatively constant body temperature (T_b_) throughout the year (Cooper and Swanson, 1994; Liknes and Swanson, 1996; Marsh and Dawson, 1989; Petit et al., 2013; Swanson, 1990; Swanson and Olmstead, 1999). In the winter, birds can theoretically increase their shivering thermogenesis by enhancing a variety of subordinate traits (see Swanson, 2010 for a review). These flexible modifications fall within four broad levels of physiological organization related to aerobic performance: (1) the size and structure of thermogenic muscle; (2) the supply of metabolic substrate and (3) oxygen to and within the muscle; and (4) the muscle’s cellular aerobic capacity. Each level is, in turn, composed of multiple traits for which there is evidence for avian seasonal acclimatization and/or cold acclimation (Figure 1). Many of these potential modifications may be accompanied by concomitant growth in maintenance costs. Indeed, basal metabolic rate also increases in the cold for many birds (McKechnie, 2008; Weathers and Caccamise, 1978), perhaps as a byproduct of other physiological changes (Swanson, 1991; Swanson, 2010). Failure to achieve adequate thermogenic capacity can have dramatic consequences for endothermic fitness (Hayes and O’Connor, 1999; Petit et al., 2017) such that thermogenic flexibility mediates a balance between thermoregulation and its associated energetic costs in response to changing climatic selective pressures (Swanson, 2010). Thus, thermogenic flexibility may profoundly influence endothermic physiological adaptation to temperate climates (Swanson and Garland, Jr., 2009).

**Figure 1.**
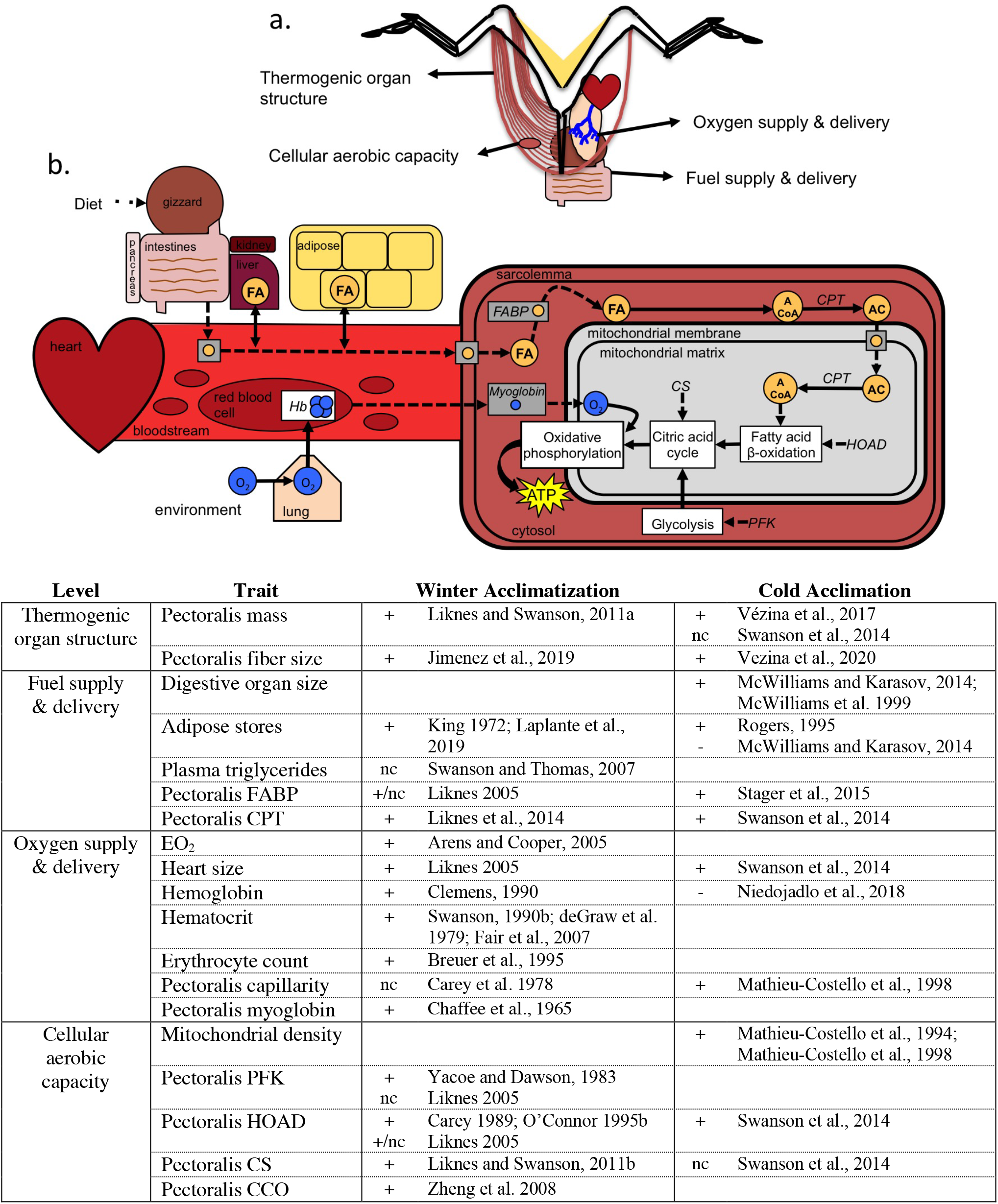
(a) Schematic of potential physiological adjustments to enhance thermogenic capacity. (B) Detail of the fuel and oxygen supply pathways as they feed into cellular aerobic metabolism. Modified from Stager et al. (2015). (c) Evidence for winter acclimatization and cold acclimation in small birds for each trait from the literature. Increased (+), decreased (−), or no change (nc).

Despite evidence for modifications to each of these subordinate traits across species, though, often only a few traits are measured in any given study (but see Vézina et al., 2017). In order to understand the relative contribution of these subordinate traits to avian thermogenic flexibility, they must instead be evaluated simultaneously. To address this knowledge gap, we conducted a large acclimation experiment aimed at investigating the mechanisms underlying thermogenic flexibility in the Dark-eyed Junco (*Junco hyemalis*). Juncos overwinter at high latitudes across North America and show increases in peak thermogenesis (the maximum metabolic rate under cold exposure; M_sum_) and cold tolerance in winter (Swanson, 1990). We exposed juncos to temperature treatments of varying duration (from one to nine weeks) and previously reported that cold-acclimated juncos increased their M_sum_ and the ability to maintain their T_b_ during acute cold exposure (Stager et al., 2020). Here we add 20 additional organ- and tissue-level phenotypes for these same individuals to explore the degree to which flexibility in subordinate physiological traits contributed to thermogenic flexibility. Specifically, we assayed body composition, organ size, muscle histology, blood parameters, and mitochondrial enzyme activities of the pectoralis representing indices of all four of the levels of physiological organization laid out above. We predicted that if avian thermogenic flexibility is a symmorphotic response, birds should make changes to traits across all four physiological levels concurrently. If, instead, control of this flexible response is concentrated in key parts of the physiological cascade, we expected birds to make changes to only a subset of traits. This comprehensive line of inquiry allows us to characterize the avian thermogenic response to cold in unprecedented detail and assess the relative contributions of component traits to whole-organism performance.

## METHODS

### Acclimations treatments

The methods for capture, acclimation, and metabolic assays have been previously described (Stager et al., 2020). Briefly, we captured adult juncos near the end of the breeding season in Missoula County, Montana, USA (~47.0°N, −113.4°W) in 2016 and 2017. We transferred birds to husbandry facilities at the University of Montana and housed them individually in common conditions for 42 days (18°C, 10h light: 14h dark), which we refer to as the “adjustment period.” We verified that breeding traits (brood patches and cloacal protuberances) were not present after this six-week adjustment period. For five additional males not included in the study, we confirmed by dissection that testes had regressed before the acclimations began.

After the adjustment period, we randomly assigned individuals to one of ten experimental groups: we subjected them to one of two temperature treatments, *Cold* (−8°C) or *Control* (18°C), lasting 1, 2, 3, 6, or 9 weeks in duration. Photoperiod was maintained at a constant 10L: 14D in all treatments and food and water were supplied ad libitum. We did not repeat the *Week 9* treatments in 2017, thus final samples sizes are *n* = 12 per treatment, except *n*_*Control_1*_ =11, *n*_*Control_9*_ = *6*, *n*_*Cold_9*_ = 5.

### Metabolic assays

We assayed M_sum_ and resting metabolic rate (RMR) using open-flow respirometry at three sampling points: capture, before and after acclimations (referred to as pre- and post-acclimation, respectively). Data for pre- and post-acclimation measures are published in (Stager et al., 2020). We assayed RMR in the evening on the day of capture and M_sum_ the following morning using methods identical to those detailed for pre-acclimation assays (see Stager et al., 2020 for details). In brief, birds were placed in a modified 1-L plastic Nalgene container for metabolic trails. RMR trials were conducted in the dark at 27°C over 3 h with ambient, dried air pumped in at 500 ml/min. Three individuals were assayed at once such that we rotated among individuals every 20 min for recording. M_sum_ trials were conducted at −5°C for ≤ 1 hr using heliox (21% O_2_, 79% He) at 750 ml/min. For both trials, the outflow from the animal’s chamber was dried, scrubbed of CO_2_, and dried again before the O_2_ concentration was quantified using a Foxbox (Sable Systems). We quantified O_2_ consumption according to Lighton (2008) using custom scripts (https://github.com/Mstager/batch_processing_Expedata_files). We defined RMR as the lowest O_2_ consumption averaged over a 10-min period and M_sum_ as the highest O_2_ consumption averaged over a 5-min period.

### Body composition assays

Body mass (M_b_) was quantified before each metabolic measurement began. In 2016, we additionally measured M_b_ on two dates during the adjustment period (roughly one week and two weeks after capture) to assess mass gain as birds acclimated to captivity. As a structural index of size, we measured the length of both tarsi (± mm) post hoc and calculated the mean tarsus length for each individual. One individual was missing its left foot at capture; thus, the right tarsus was used as the mean.

Immediately before each M_sum_ trial, we also assayed body composition using quantitative magnetic resonance (EchoMRI Whole Body Composition Analyzer). This allows for rapid quantification of fat, lean, and water masses without sedation (Guglielmo, 2010). We quantified body composition three times for each individual—at capture, before and after acclimation— which allows us to use lean mass as a proxy for organ and muscle masses during the first two time points when destructive sampling was not possible. We calibrated the instrument daily before measurements began. We also assayed an oil standard at the beginning and end of a day’s measurements. We used the variation in the standard measures across the day’s two time points to calculate a daily rate of drift for each fat, lean, and water masses. Individual measures were then linearly corrected using this rate of drift (slope) and the initial deviation from the standard measure (intercept). We report fat mass, lean mass, free water, and total water in grams.

### Blood parameters

Directly following the pre- and post-acclimation M_sum_ trials, we extracted blood from the brachial vein to quantify blood O_2_ parameters. We first collected 10 μl of whole blood in a cuvette to assay hemoglobin concentration (g/dL) using a Hemocue Hb 201+ analyzer. To quantify hematocrit levels, we collected ~50 μl of blood, centrifuged it for 5 min, and measured the proportion of packed red blood cells to total blood volume.

Post-acclimation, we collected an additional blood sample from the jugular vein. To quantify, erythrocyte number we mixed 10 μl whole blood with 1990 μl of 0.85% saline and later imaged 10 μl of solution on a Neubauer hemocytometer. We randomly selected one of the twenty-five central grid cells (0.04 mm^2^) in which to count erythrocytes. Samples that were not imaged within 5 days of blood collection were removed from analysis due to degradation of the sample. We centrifuged the remaining blood sample to separate the red blood cells, then pipetted off the plasma, flash-froze and stored it at −80°C for future assays.

As an index of fat mobilization capacity, we quantified plasma lipid metabolites by endpoint assay on a microplate spectrophotometer at a later date. Assays were run according to Guglielmo et al. (2002a) in 400 μl flat-bottom 96 well polystyrene microplates. We thawed plasma and diluted samples three-fold with 0.9% NaCl. We first measured free glycerol concentration (5 μl plasma, 240 μl free glycerol Sigma reagent A) at 37°C and A540. We then added 60 μl triglyceride (Sigma reagent B) and read absorbance at the same spectrophotometer conditions to quantify total triglyceride concentrations. Samples were run in duplicate and standard curves were included for each plate. Intra-assay and inter-assay coefficients of variation were 0.35 and 0.34 for total triglycerides and 0.24 and 0.36 for glycerol, respectively. True triglyceride concentration (TRIG) was calculated as total triglyceride minus glycerol (mmol L^−1^).

### Organ masses

At the end of the acclimation treatments, immediately following the final M_sum_ trial and blood extraction, we euthanized individuals using cervical dislocation. We excised the left pectoralis for enzyme assays and the right pectoralis for histological purposes (see below). We weighed organs with a 0.0001 g precision balance (Mettler Toledo ME104). We excised the heart, removed major vessels, fat, and blood before weighing it, and similarly preserved it for histology. We harvested the liver, right kidney, and lungs, trimmed fat, blotted blood on the surface, weighed each (wet mass), and then dried them at 60°C for 48 h before quantifying dry mass. Lungs were not completely exsanguinated, thus blood content likely contributed to mass. Right and left lung masses did not differ (t-test: t = −0.67, df = 206, p = 0.50) and are reported as total lung mass. In 2017, we additionally harvested the gizzard (proventriculus removed), intestines (from gizzard to cloaca; small and large combined), spleen, and pancreas in the same way. Gonads were regressed in all cases and were not weighed. We report wet mass for heart, and dry mass for all other organs (spleen not shown in text).

Due to the difficulty of quantifying total muscle mass directly, we approximated muscle size with data from 2017 individuals. First, we totaled all wet organ masses and subtracted this value from lean mass. We did this multiplying kidney mass by two and using a proportionally constant estimate of brain mass from the literature based on an individual’s mass at capture because we did not expect brain mass to change with acclimation. We used the remaining value as an index of wet muscle mass and assumed 75% water content to arrive at a rough estimate of dry muscle mass. This estimate includes other organs not measured here that may have responded to our acclimation treatments (e.g., esophagus, crop, proventriculus). To validate this measure, we separately estimated the water content of muscle by calculating water composition for each organ (wet minus dry masses) and subtracting these values, as well as the mass attributed to free water, water in fat, and water in other tissues (i.e., bones, skin, feathers) from the total water mass for each individual. To do this, we approximated brain mass as before, and estimated that brain and heart (for which we did not quantify dry mass) were composed of 77% and 75% water, respectively (Graber and Graber, 1965; Hughes, 1974). We also assumed that adipose stores were composed of 10% water and that M_b_ not assigned to lean, fat, or free water could be attributed to bones, skin, and feathers, for which we estimated 20% water content. Though rough approximations, these independent estimates of dry muscle mass and water content of the muscle are strongly correlated (Pearson’s correlation: *r* = 0.84, *p* = 5.8 × 10^−14^).

### Muscle histology

The pectoralis is the principle muscle used for shivering in small birds (Yacoe and Dawson, 1983). In 2017, we excised the middle section of the right pectoralis, coated it with embedding medium (OCT compound), froze it in a bath of isopentane, and stored the sample at −80°C until sectioning. We sectioned pectoralis tissue (10 μm) transverse to muscle fiber length at −20°C using a Leica CM1950 Cryostat. We mounted sections on poly-L-lysine–coated slides, air-dried and stored them at −80°C until staining occurred. To identify capillaries, we stained for alkaline phosphatase activity. We first incubated slides at room temperature for ~2 h then fixed them in acetone for 5 min and allowed them to air dry. We stained slides in assay buffer (1.0 mM nitroblue tetrazolium, 0.5 mM 5-bromo-4-chloro-3-in- doxyl phosphate, 28 mM NaBO_2_, and 7 mM MgSO_4_) at pH 9.3 for 1 h. We imaged muscle sections using light microscopy and used stereological quantification methods to make unbiased measurements (Weibel 1979; Egginton 1990). For a randomly selected subset (200 mm^2^) of the image, we then quantified capillary number relative to muscle fiber count and capillary density (per mm^2^). We analyzed three regions for each sample to account for heterogeneity across the tissue.

### Enzyme assays

Upon excision, we flash froze the left pectoralis in liquid nitrogen, stored it at −80°C, and later used it to quantify activities of carnitine palmitoyl transferase (CPT; an indicator of fatty acid transport into the mitochondrial membrane), beta hydroxyacyl Co-A dehydrogenase (HOAD; an indicator of fatty acid oxidation capacity), and citrate synthase (CS; an indicator of maximal cellular metabolic intensity) according to Guglielmo et al., (2002b). We combined 100 mg frozen pectoralis tissue with 9 volumes ice-cold homogenization buffer (20 mM Na_2_HPO_4_, 0.5 mM EDTA, 0.2% fatty acid-free BSA, 0.1% Triton X-100, and 50% glycerol at pH 7.4). We homogenized tissues for 3 min at high speed using a Qiagen TissueLyser with adapter sets cooled to −20°C. We further diluted crude muscle homogenates to 1:100 with homogenization buffer, divided samples, and stored aliquots at −80°C until assays were performed. Maximal enzyme activities were quantified using a microplate spectrophotometer. All assays were performed in duplicate, in 400 μl flat-bottom 96 well polystyrene microplates at 39°C, with a reaction volume of 200 μl. Assay conditions were: 50 mM Tris buffer pH 8.56, 7.5 mM carnitine, 0.035 mM palmitoyl-CoA, 0.15 mM DTNB, and 20 μl diluted homogenate for CPT; 50 mM imidazole pH 7.96, 1 mM EDTA, 0.1 mM aceto-acetyl-CoA, 0.2 mM NADH, and 20 μl diluted homogenate for HOAD; and 50 mM Tris buffer pH 8.56, 0.75 mM oxaloacetic acid, 0.10 mM acetyl-CoA, 0.15 mM 5,5-dithiobis-(2-nitrobenzoic acid) (DTNB), and 2 μl diluted homogenate for CS. Activities (μmol•min^−1^) were calculated from A412 (ε = 13.6) for CS and CPT and from A340 (ε = 6.22) for HOAD. Week 9 individuals were not included for CPT and CS assays.

### Statistical analyses

We performed all analyses in the statistical environment R (R Core Team, 2018). We performed analysis of variance tests to verify that the ten treatment groups did not differ in trait values either at capture or before acclimation (Tables S1). To quantify the rate of mass gain across the adjustment period, we employed the repeated measures of M_b_ obtained in 2016 in a linear mixed model with days in captivity as a fixed effect and individual as a random effect. We used pairwise t-tests to assess changes in body composition that occurred between capture and the pre-acclimation assays.

To compare the relative degree of change among phenotypic traits in response to temperature acclimation, we first standardized each phenotypic variable (by subtracting the mean and dividing by two standard deviations) using the package *arm* (Gelman, 2008). We tested for effects of Treatment, Duration, and their interaction on phenotypic measures using linear models. In all cases, Treatment × Duration terms were not significant (Table S2) and thus models without the interaction are presented in the text. We also used linear models to test for an effect of *Year* on phenotypic measures that were repeated in both years of the study. We established significance after Bonferroni corrections for multiple testing.

We tested for pairwise associations between all phenotypic traits for a given sampling period with Pearson’s correlation tests. In order to determine the relative influence of subordinate phenotypes on M_sum_, we utilized the variation in traits exhibited across temperature treatments post-acclimation and performed regressions of standardized trait values on M_sum_. Rather than including all possible traits, we used only those identified with Pearson’s correlations to be associated with post-acclimation M_sum_. Including single terms in each model allowed us to maximize sample sizes for each trait and avoid complications associated with combining terms, like lean mass, which is a composite trait and would therefore be redundant to measures of muscle and organ masses.

## RESULTS

### At capture

At capture, 10 of 15 pairwise trait combinations (67%) showed correlations. Juncos that were structurally larger were also heavier and carried more lean mass (Figure 2a), but all birds had very little fat (Table S3). Differences did not exist in body size or composition between years, yet individuals exhibited slightly higher metabolic rates in 2017 (Table S4). M_sum_ positively correlated with M_b_, lean mass, and RMR (Figure 2a).

**Figure 2.**
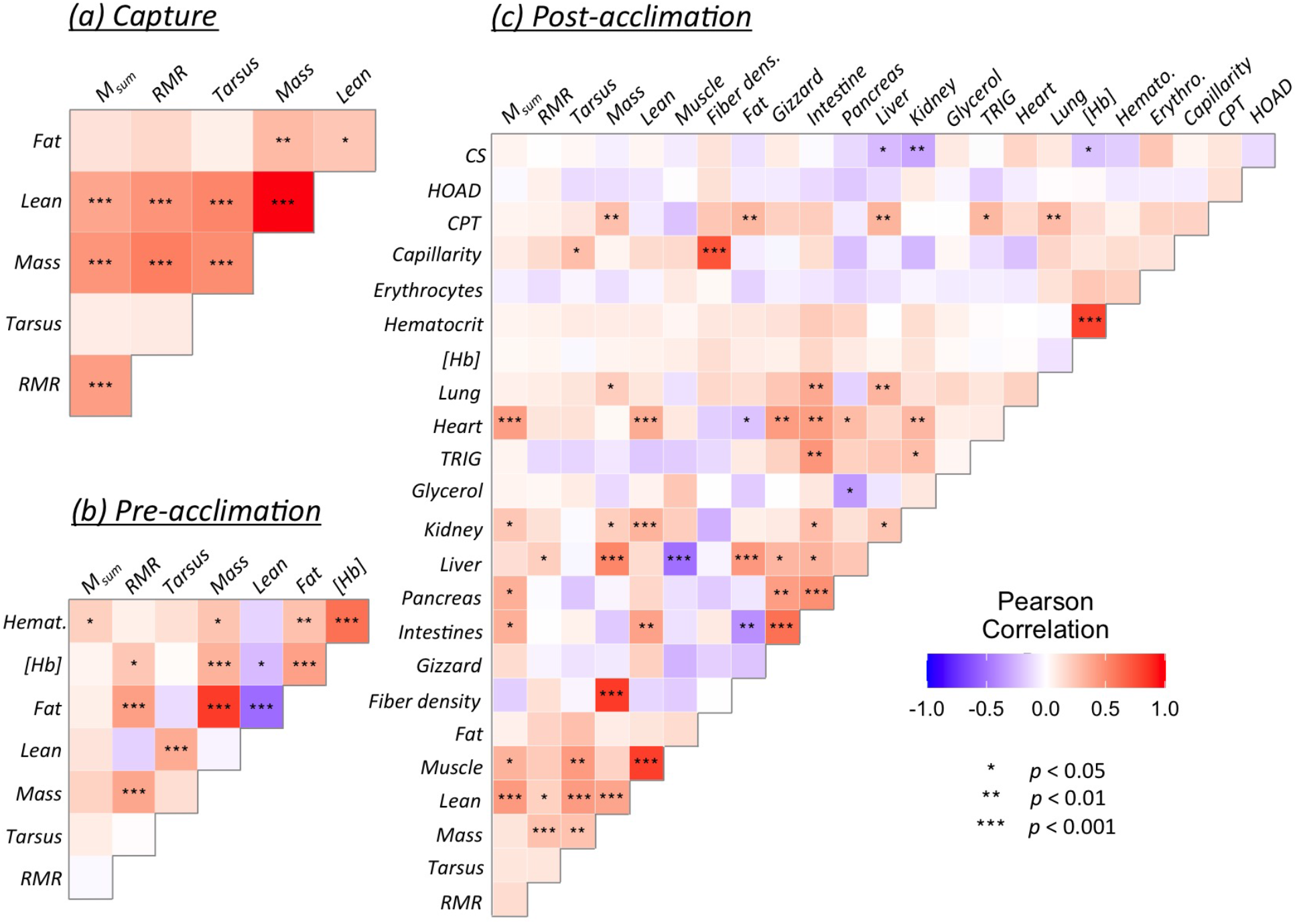
Pairwise trait correlations at (a) capture, (b) before acclimation, and (c) after acclimation. Colors correspond to Pearson’s correlation coefficients (positive = red; negative = blue); asterisks indicate significance. Underlying values shown in Tables S5–S7.

### Prior to acclimation

Our six-week adjustment period successfully reduced variation in M_sum_ among individuals (var_*Capture*_ = 2.11 vs. var_Pre_ = 0.36 ml O_2_•min^−1^). Juncos rapidly increased M_b_ over this time (Table S3), with birds gaining 0.10 g per day in 2016. Most of this mass gain can be attributed to growth in adipose stores, though birds did increase lean mass to a lesser degree (Table S3). Individuals gained more M_b_ — particularly fat mass — during the six-week adjustment period in 2017 than in 2016 (Table S4). Importantly, treatment groups did not differ at capture or before acclimations for any of the phenotypic traits assayed (Table 1; Table S1).

**Table 1.**
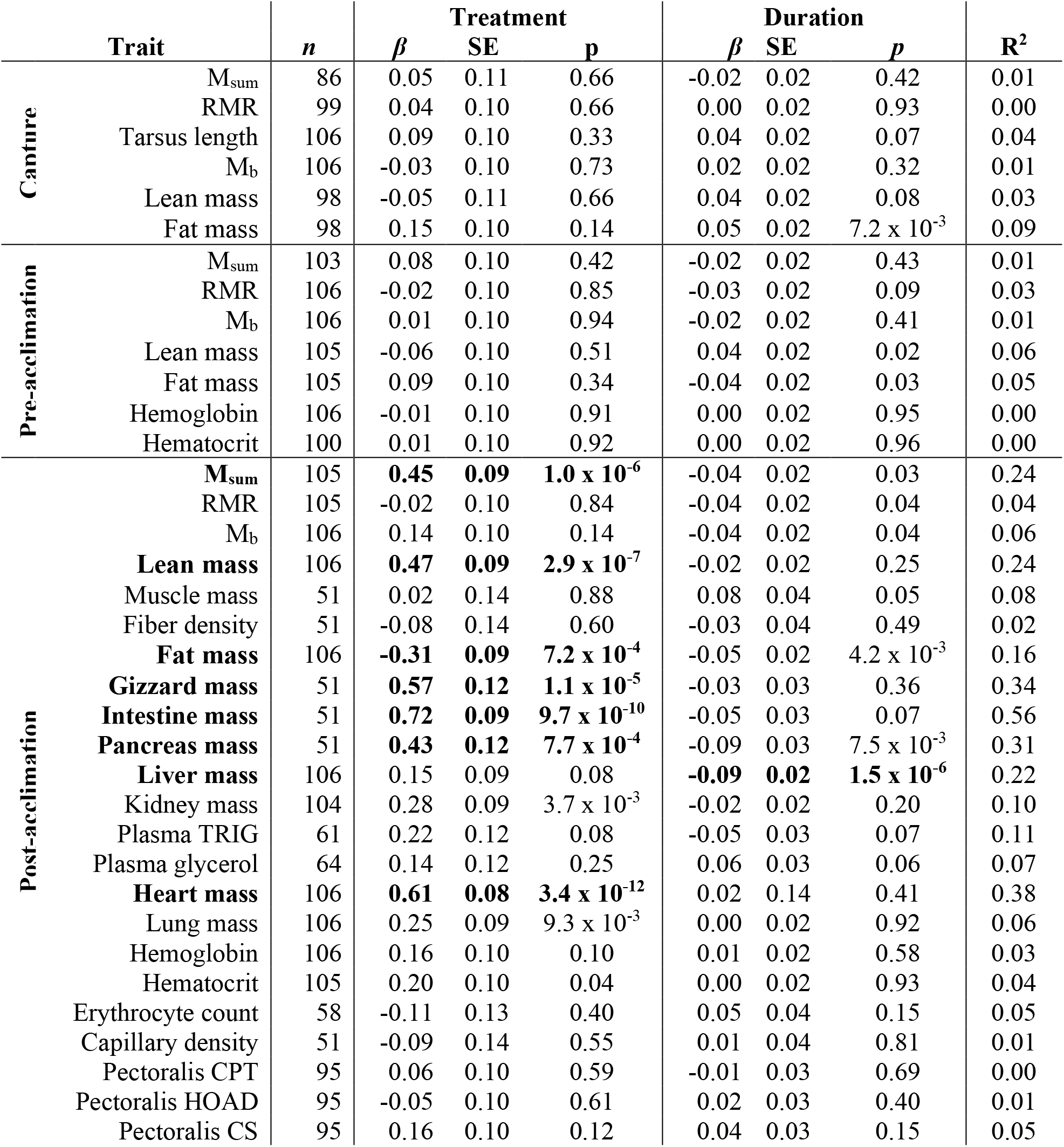
Linear effects of *Cold* treatment and *Duration* on standardized values of phenotypic traits. Significant effects after correction for 36 tests (p < 0.0014) bolded. Metabolic rates, body mass, and tarsus lengths for pre- and post-acclimation from Stager et al. (2020).

Immediately prior to acclimation, 13 of 28 pairwise trait combinations (46%) exhibited correlations (Figure 2b). Only 3 of these associations were present at capture. M_sum_ correlated positively with hematocrit alone.

### After acclimation

Several trait values were modified in response to cold acclimation. Although Mb did not vary among treatments, juncos adjusted body composition in the cold (Table 1). Cold-acclimated individuals exhibited 0.73 g more lean mass and 0.92 g less fat mass compared to *Control* individuals. This difference in lean mass can, in part, be attributed to growth of the digestive tract in *Cold* birds, which increased the size of their gizzard, intestines, and pancreas by 39%, 49%, and 28% respectively relative to *Control* birds. Cold-acclimated juncos additionally enlarged the size of their heart by 15% compared with *Control* individuals. Both lung mass and kidney mass increased in the cold, but these trends were not significant after correction for multiple testing. Liver mass, which decreased over time in both temperature treatments, was the only trait to show a significant effect of treatment duration. In contrast, muscle, blood, and enzymatic parameters exhibited little flexibility among treatments.

After acclimation, 52 of 276 pairwise trait combinations (19%) exhibited correlations (Figure 2c). Of these associations, 7 were also observed at capture and 4 were observed before acclimation. Only 3 associations were common to all three contexts: RMR and M_b_; fat mass and M_b_; and lean mass and tarsus length. Six traits correlated positively with M_sum_ after acclimation, and most involved organ masses that had not been measured at prior sampling points. Lean and heart masses showed the strongest influence on M_sum_, exhibiting effects equal in magnitude and direction (Table 2).

**Table 2.** Effects of standardized phenotypic traits on M_sum_ post-acclimation.

|  | **Phenotypic Trait** | ***n*** | ***β*** | **SE** | ***p*** | **$R^2$** |
| --- | --- | --- | --- | --- | --- | --- |
| **Post-acclimation** | Lean mass | 105 | 1.26 | 0.26 | $3.5 \times 10^{-6}$ | 0.19 |
| | Heart mass | 105 | 1.25 | 0.27 | $1.0 \times 10^{-5}$ | 0.17 |
|  | Muscle mass | 50 | 1.11 | 0.46 | 0.02 | 0.11 |
|  | Kidney mass | 103 | 0.71 | 0.29 | 0.02 | 0.06 |
|  | Intestine mass | 50 | 1.14 | 0.46 | 0.02 | 0.12 |
|  | Pancreas mass | 50 | 1.15 | 0.46 | 0.02 | 0.12 |

## DISCUSSION

Phenotypic flexibility allows individuals to change trait values in order to match their phenotypes with fluctuations in environmental conditions. Although many whole-organism phenotypes are composed of a complex network of subordinate traits, the coordinated ways in which these lower-level traits respond to environmental challenges and contribute to phenotypic flexibility has been little explored. We previously demonstrated that adult Dark-eyed Juncos increased their thermogenic capacity (M_sum_) in response to synthetic temperature cues, and that this increase corresponded with an enhanced ability to maintain T_b_ in the cold (Stager et al., 2020). Here we add measures of 20 additional sub-organismal, physiological traits for the same individuals, several of which were measured repeatedly in the same individuals, providing a rich dataset for exploring the mechanistic basis of phenotypic flexibility. We show that the relationships among a number of these traits varied as the environmental context changed. Moreover, variation in M_sum_ in response to temperature acclimation was correlated with only a handful of subordinate phenotypes. Our results thus indicate that avian thermogenic flexibility is not a symmorphotic response, but rather that adjustments to thermogenic flexibility are concentrated in a few subordinate traits. If this is a general feature of complex traits, it suggests that the evolution of phenotypic flexibility in a single component part could impart flexibility for the system as a whole, thereby enabling simple and reversible modifications to significantly impact whole-organism performance in response to environmental change.

### Phenotypic responses to cold

We found that in response to very cold temperatures, juncos increased lean mass by enlarging the size of several major organs and simultaneously decreased adipose stores relative to *Control* birds. These traits changed rapidly and plateaued within the first week of cold exposure such that increased duration of the temperature treatment had little effect on trait values. Juncos were thus able to respond on shorter timescales to a significant environmental stressor than had previously been appreciated (see also <u>Stager et al., 2020</u>).

Intriguingly though, many other traits that have been previously implicated in avian thermogenic flexibility remained unchanged. For example, we hypothesized that increased thermogenic capacity might be achieved by augmenting the fuels supporting aerobic metabolism — either directly from food processing or from reserves. While we cannot address the former, counter to the latter idea, juncos had lower adipose depots in the cold, similar to cold-acclimated White-throated Sparrows (*Zonotrichia albicollis*; McWilliams and Karasov, 2014). This change in body composition could result from cold-acclimated individuals burning fat faster than they were able to store it. Accordingly, we observed that juncos gained, on average, only 0.10 g of M_b_ per day at 18°C during the adjustment period, which is likely not enough to overcome rates of overnight mass loss at cold temperatures (e.g., Ketterson and Nolan, 1978).

Consequently, with fat stores likely being quickly diminished, *Cold* birds may have instead maintained their fuel supplies by increasing food consumption. Many birds accompany higher food intake with growth to their digestive track, which allows individuals to process larger food quantities more quickly without losses to digestive efficiency (McWilliams and Karasov, 2001). In support of this, although we did not quantify food intake, we did find that birds increased the size of their gizzard, intestine, and pancreas within the first week of cold acclimation. Likewise, White-throated Sparrows grew their intestines within 2-12 d of cold exposure (−20°C), and their larger digestive tracks facilitated greater digestive capacity and increased feeding rates in the absence of reciprocal changes in nutrient uptake per unit of intestine (McWilliams and Karasov, 2014). If true of juncos as well, at high rates of energy use, this could enable them to efficiently use digestive products without spending energy to convert fuels to/from stored adipose. However, we did not observe increases in fat transporters either in the blood or within the muscle. The fact that all traits did not change suggests that many traits may harbor spare capacity (*sensu* McWilliams and Karasov, 2001) such that they can accommodate larger demands without significantly adjusting their trait value.

Ultimately, our treatments lasted up to two months and birds were exposed to a fairly extreme temperature stimulus such that the phenotypic responses shown here are not likely to have been constrained by time or insufficient severity of the cue. Additionally, junco M_sum_ plateaued within one week of cold acclimation (Stager et al., 2020). As a result, any discrepancies between our findings and those shown in wild birds may follow from the fact that winter acclimatization likely involves the combination of several environmental cues. We focused on temperature specifically because previous work has shown that junco M_sum_ responds to variation in temperature rather than photoperiod (Swanson et al., 2014). This had the benefit of allowing us to isolate the phenotypic responses that underlie thermogenic performance in order to decompose this complex trait. Nonetheless, it means that we may have missed certain hormonal changes and subsequent physiological responses that are likely tied to photoperiod or variation in resource abundance and availability, and thus associated with the “winter phenotype.” We cannot therefore discount the fact that a different environmental cue — or several coinciding cues — may induce maximal output at every level through the regulation of trait changes (i.e., symmorphosis).

### Variation in trait associations across time

Even though our acclimation treatment targeted responses to cold alone, an unintended outcome of our experimental setup is that several environmental variables changed throughout the course of the investigation as a whole. For instance, juncos were nearing the end of their breeding season when they were captured, which is typically considered a “lean” time of the annual cycle. In addition to defending territories and provisioning young, they were likely contending with variation in temperatures, precipitation, food availability, and predation pressures in the wild. These stressors are reflected in the poor body condition of our birds at capture. In contrast, during the adjustment period in the lab environment, breeding traits quickly regressed following exposure to an artificially short photoperiod, and birds were housed individually with ad libitum food under mild temperatures (albeit, outside of their thermoneutral zone). These conditions therefore represented a more benign environment than that experienced by wild juncos at this time of year, and the standard conditions removed inter-individual variation. When we next induced temperature changes, we did so in the absence of variation in photoperiod or food availability, after birds had already adjusted to captivity. The phenotypic responses that we observed during this period are therefore reflective of the isolated effect of cold temperatures. Thus, because birds must respond to changes in their environment across many different axes in the wild, these three contexts let us explore how consistently traits may be associated.

In total, we quantified 253 pairwise trait correlations among individuals, 28 of which we measured two or more times per individual (e.g., at capture, before acclimation, and after acclimation). Many of these relationships varied in either strength or direction across the three sampling periods. For example, fat and lean masses, which correlated positively at capture, correlated negatively after six weeks of captivity. All individuals were thus capable of storing adipose in this setting. At this same time point, though, other trait correlations that existed at the time of capture were absent, perhaps due to the reduced phenotypic variation following the adjustment period. Ultimately, of the original trait associations exhibited at capture, 70% did not persist across the subsequent sampling points.

Notably, the traits that correlated with M_sum_ also changed across time, such that ratios between subordinate traits and aerobic performance were not invariant to perturbation, as would be predicted by symmorphosis. Lean mass correlated with M_sum_ both at capture and after, but not before, acclimation. Meanwhile, RMR and M_b_ correlated with M_sum_ only at capture and not once birds had adjusted to captivity. Collectively, these results point to the importance of environmental context in evaluating phenotypic contributions to performance and, more broadly, imply that relationships between flexible phenotypic and performance traits — which are often used as indices of fitness — may change across time.

### Symmorphosis?

Previous work has indicated that symmorphosis may be generally applicable to the limits of avian aerobic performance (Seymour et al., 2008; Suarez, 1998; Swanson, 2010). However, because most studies focus on the contribution of oxygen transport alone, they could also be interpreted as demonstrating that correlations exist across only *some* of the physiological pathways associated with aerobic performance (Swanson, 2010). In comparison, we did not observe correlations among the many parameters quantified within the oxygen supply pathway, but we did see associations between several traits related to fuel transport. In addition to correlations among many of the digestive organs, these organs positively correlated with heart mass, and plasma triglyceride concentrations positively correlated with intestinal mass and with CPT activity, as well. Together this indicates that higher digestive capacity was likely met with higher fuel transport capacity.

In its strictest sense, though, symmorphosis predicts that all components within a system should change in concert such that no one element is operating in excess (Weibel et al., 1991). We did not find support for this hypothesis in that juncos achieved higher thermogenic capacity in the cold without simultaneously adjusting each subordinate trait. Juncos instead enhanced M_sum_ by concurrently modifying five traits that fall within three levels of biological organization (Figure 1) — including the masses of the muscle, certain digestive organs, and the heart; however, they did not alter lower-level indices of cellular aerobic capacity that we measured here.

Growing larger organs may seem like a costly and time-consuming investment for a small bird to make if an alternative possibility is to increase the expression of key metabolic enzymes. Though we did not quantify their cost of production, surprisingly, none of these organ sizes correlated with RMR, suggesting that larger organs were not associated with higher maintenance costs as predicted (e.g., Chappell et al., 1999; Vézina et al., 2017). Moreover, sizeable growth in these traits was achieved within one week of cold exposure indicating that these modifications are induced on seemingly short time scales rather than preemptive to seasonal temperature changes. Thus, in order to fully understand the costs of trait production as they relate to reversible modification, de-acclimation studies are also needed.

Of any single trait, heart mass exhibited the largest effect on M_sum_. Unfortunately, though, we did not quantify as many traits in the first year of the study as we did in the second. This may have reduced our power to detect associations among traits and appropriately assess their relative influence on M_sum_. For instance, the strong influence of lean mass (measured in both years) on M_sum_, in combination with the strong correlations between lean mass and muscle (*r* = 0.82) and intestinal (*r* =0.40) masses in 2017, is suggestive that these phenotypes likely influenced M_sum_ in 2016, as well. If so, their effects may have outweighed that of heart mass across all individuals. This would not be surprising as the potential benefit of larger muscles to facilitate shivering is clear, and the advantage of larger intestine, pancreas, and kidney masses likely derives from a greater digestive and excretory capacity to fuel aerobic performance, as discussed above. However, cardiac function is involved in both the fuel and oxygen supply pathways, suggesting that enhancements to this one component could have dual benefits. Increased heart size may therefore be an especially efficient way to increase flux across multiple parts of the physiological network.

## Conclusions

Understanding how organisms flexibly alter physiological responses can help us understand their capacity to cope with a changing environment (Stillman, 2003). Taken together, our results indicate that flexibility in a whole-organism performance phenotype can be modified quickly by altering a handful of underlying traits of large effect. If this is a generalizable feature of phenotypic flexibility, it may help explain its ubiquity across many morphological, physiological, and behavioral traits. We thus urge future studies to continue exploring how flexibility in performance traits is achieved and to develop a cost-benefit framework that can help put into context why some traits are flexible, while others are not.

## Acknowledgements

We are especially thankful to Luke Wilde for help quantifying histological traits. We also thank Nathan Senner, Phred Benham, Ryan Mahar, and Nicholas Sly for their help in catching birds. We thank Chris Guglielmo for sharing his plasma metabolite assay protocols; Jesse Loewecke, Catie Ivy, and Lou Herritt for histology assistance; Frank Moss for logistical assistance; Hailey Bunker for help with husbandry; Alex Gerson and Francois Vézina for helpful advice after the initial year of data collection; as well as Keely Corder, Ryan Mahar, Rena Schweizer, Jon Velotta, and Cole Wolf for feedback on an earlier draft of this manuscript.

## Ethics

All procedures were approved by the University of Montana Animal Care Committee (Protocol 010-16ZCDBS-020916). Birds were collected with permission from Montana Fish Wildlife & Parks (permits 2016-013 and 2017-067-W, issued to M.S.) and the US Fish & Wildlife Service (permit MB84376B-1 to M.S.).

## Author Contributions

M.S. and Z.A.C. conceived of the study; M.S. performed acclimations, data collection, and analyses, and drafted the manuscript; Z.A.C. contributed edits to the manuscript.

---

**Table S1.**
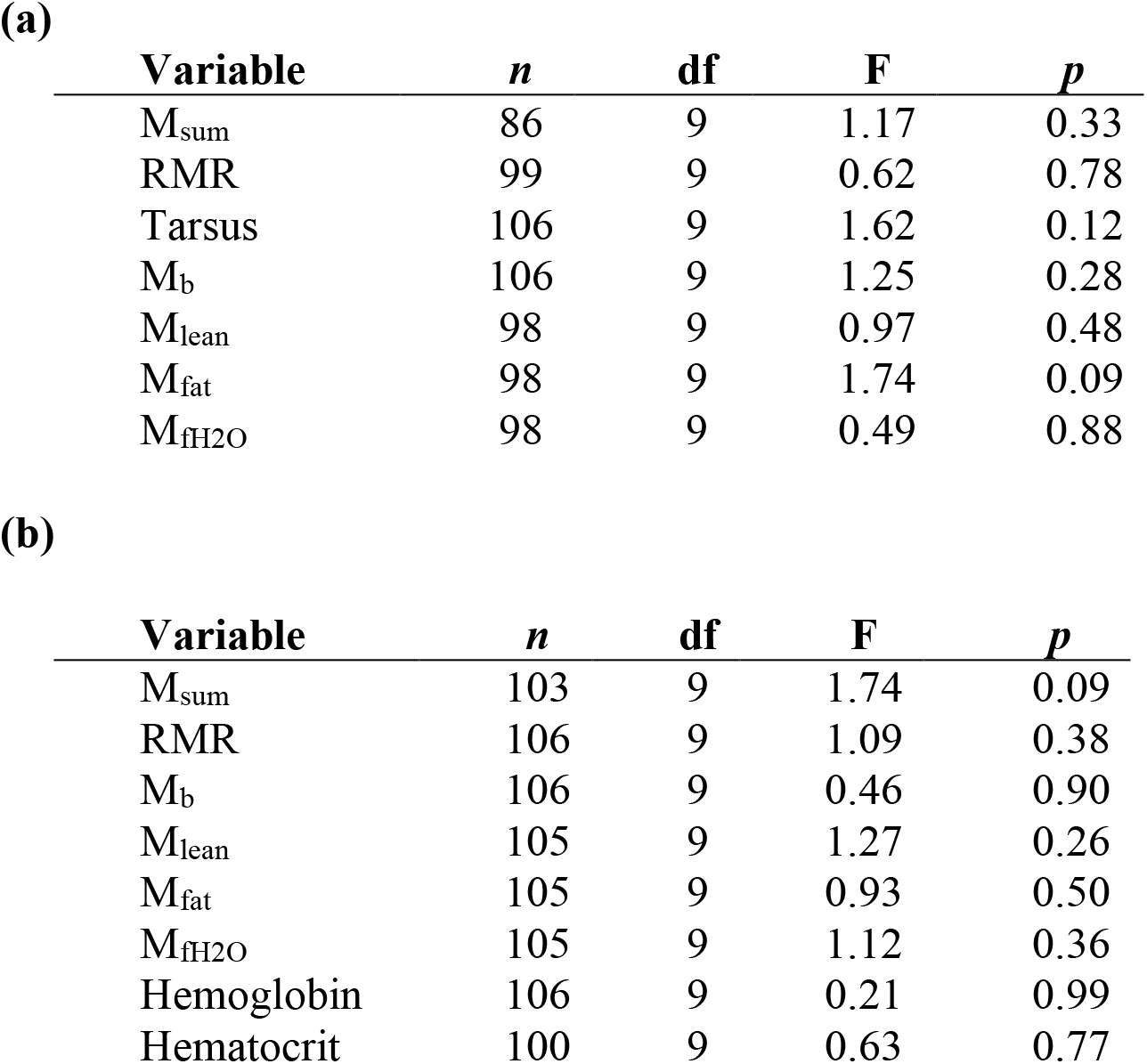
Results from ANOVAs to determine if the ten treatment groups differed in physiological parameters at (a) capture and (b) before acclimation treatments.

**Table S2.** Linear effects of *Cold* Treatment, *Duration*, and their interaction on standardized phenotypic traits. Body mass, tarsus lengths, and metabolic rates for pre- and post-acclimation from Stager et al. (2020). Bolded significant effects after Bonferroni correction for 36 models (p < 0.0014).

|  | Trait | *n* | Treatment |  |  | Duration |  |  | Treat x Duration |  |  |
| --- | --- | --- | --- | --- | --- | --- | --- | --- | --- | --- | --- |
| | | | $\beta$ | SE | p | $\beta$ | SE | p | $\beta$ | SE | p |
| Canture | RMR | 99 | 0.13 | 0.18 | 0.48 | 0.01 | 0.03 | 0.76 | -0.02 | 0.04 | 0.58 |
|  | M<sub>sum</sub> | 86 | -0.15 | 0.20 | 0.46 | 0.04 | 0.03 | 0.17 | 0.05 | 0.05 | 0.24 |
|  | Tarsus Length | 106 | 0.06 | 0.17 | 0.73 | 0.03 | 0.03 | 0.25 | 0.01 | 0.04 | 0.79 |
|  | M<sub>b</sub> | 106 | 0.06 | 0.17 | 0.72 | 0.03 | 0.03 | 0.24 | -0.03 | 0.04 | 0.50 |
|  | M<sub>lean</sub> | 98 | -0.15 | 0.18 | 0.41 | 0.02 | 0.03 | 0.38 | 0.03 | 0.04 | 0.49 |
|  | M<sub>fat</sub> | 98 | 0.27 | 0.17 | 0.12 | 0.07 | 0.03 | 0.01 | -0.03 | 0.04 | 0.40 |
| Pre-acclimation | RMR | 106 | 0.08 | 0.17 | 0.65 | -0.02 | 0.03 | 0.45 | -0.03 | 0.04 | 0.49 |
|  | M<sub>sum</sub> | 103 | 0.24 | 0.18 | 0.18 | 0.00 | 0.03 | 0.88 | -0.04 | 0.04 | 0.29 |
|  | M<sub>b</sub> | 106 | -0.10 | 0.17 | 0.55 | -0.03 | 0.03 | 0.26 | 0.03 | 0.04 | 0.43 |
|  | M<sub>lean</sub> | 105 | -0.12 | 0.17 | 0.47 | 0.04 | 0.03 | 0.17 | 0.02 | 0.04 | 0.67 |
|  | M<sub>fat</sub> | 105 | 0.12 | 0.17 | 0.50 | -0.04 | 0.03 | 0.15 | -0.01 | 0.04 | 0.87 |
|  | Hemoglobin | 106 | -0.05 | 0.17 | 0.76 | -0.01 | 0.03 | 0.80 | 0.01 | 0.04 | 0.76 |
|  | Hematocrit | 100 | -0.03 | 0.18 | 0.87 | 0.00 | 0.03 | 0.88 | 0.01 | 0.04 | 0.79 |
| Post-acclimation | RMR | 105 | -0.24 | 0.17 | 0.16 | -0.07 | 0.03 | 0.01 | 0.06 | 0.04 | 0.11 |
| | M<sub>sum</sub> | 105 | 0.44 | 0.15 | $4.0 \times 10^{-3}$ | -0.04 | 0.02 | 0.11 | 0.00 | 0.03 | 0.98 |
|  | M<sub>b</sub> | 106 | -0.01 | 0.17 | 0.95 | -0.06 | 0.03 | 0.03 | 0.04 | 0.04 | 0.27 |
|  | M<sub>lean</sub> | 106 | 0.33 | 0.15 | 0.03 | -0.04 | 0.02 | 0.11 | 0.04 | 0.03 | 0.26 |
|  | M<sub>fat</sub> | 106 | -0.36 | 0.16 | 0.02 | -0.06 | 0.02 | 0.02 | 0.01 | 0.04 | 0.69 |
|  | Hemoglobin | 106 | 0.05 | 0.17 | 0.76 | 0.00 | 0.03 | 0.88 | 0.03 | 0.04 | 0.43 |
|  | Hematocrit | 105 | 0.04 | 0.17 | 0.83 | -0.02 | 0.03 | 0.46 | 0.04 | 0.04 | 0.25 |
|  | Erythrocyte count | 58 | 0.01 | 0.27 | 0.97 | 0.07 | 0.05 | 0.16 | -0.04 | 0.07 | 0.61 |
|  | Capillary density | 51 | 0.04 | 0.28 | 0.90 | 0.03 | 0.06 | 0.59 | -0.04 | 0.08 | 0.61 |
|  | Fiber density | 51 | 0.28 | 0.27 | 0.32 | 0.03 | 0.06 | 0.56 | -0.12 | 0.08 | 0.14 |
|  | **Heart mass** | 106 | **0.46** | **0.13** | **$9.0 \times 10^{-4}$** | 0.00 | 0.02 | 0.91 | 0.04 | 0.03 | 0.18 |
|  | Lung mass | 106 | 0.35 | 0.17 | 0.04 | 0.01 | 0.03 | 0.67 | -0.03 | 0.04 | 0.47 |
|  | **Liver mass** | 106 | 0.08 | 0.15 | 0.60 | **-0.10** | **0.02** | **$9.5 \times 10^{-5}$** | 0.02 | 0.03 | 0.56 |
|  | Kidney mass | 104 | -0.01 | 0.16 | 0.94 | -0.06 | 0.03 | 0.02 | 0.08 | 0.04 | 0.03 |
| | Gizzard mass | 51 | 0.65 | 0.05 | $6.0 \times 10^{-3}$ | -0.02 | 0.05 | 0.74 | -0.03 | 0.06 | 0.67 |
|  | **Intestine mass** | 51 | **0.82** | **0.18** | **$5.9 \times 10^{-5}$** | -0.03 | 0.04 | 0.40 | -0.03 | 0.05 | 0.53 |
|  | Pancreas mass | 51 | 0.54 | 0.23 | 0.02 | -0.07 | 0.05 | 0.13 | -0.04 | 0.07 | 0.57 |
|  | Muscle mass | 51 | -0.32 | 0.26 | 0.23 | 0.02 | 0.05 | 0.31 | 0.11 | 0.08 | 0.13 |
|  | CPT | 95 | -0.06 | 0.20 | 0.75 | -0.03 | 0.04 | 0.43 | 0.04 | 0.06 | 0.48 |
|  | HOAD | 95 | -0.03 | 0.20 | 0.90 | 0.03 | 0.04 | 0.48 | -0.01 | 0.06 | 0.87 |
|  | CS | 95 | 0.22 | 0.19 | 0.26 | 0.05 | 0.04 | 0.21 | -0.02 | 0.05 | 0.71 |
|  | Plasma TRIG | 61 | 0.26 | 0.23 | 0.27 | -0.05 | 0.04 | 0.20 | 0.01 | 0.06 | 0.85 |
|  | Plasma glycerol | 64 | 0.12 | 0.23 | 0.59 | 0.05 | 0.04 | 0.16 | 0.01 | 0.06 | 0.92 |

**Table S3.** Body composition (in grams) across time points. Mean ± SD and two-sample t-test results.

| **Trait** | **Capture** | **Pre-Acclimation** | **t** | **df** | ***p*** |
| --- | --- | --- | --- | --- | --- |
| M<sub>b</sub> | 17.45 ± 1.24 | 22.60 ± 1.57 | -25.2 | 193 | $<2.2 \times 10^{-16}$ |
| Lean mass | 14.03 ± 1.01 | 14.70 ± 0.93 | -4.9 | 197 | $1.7 \times 10^{-6}$ |
| Fat mass | 0.08 ± 0.11 | 3.60 ± 1.65 | -21.8 | 105 | $<2.2 \times 10^{-16}$ |
| Free water mass | 0.33 ± 0.04 | 0.32 ± 0.06 | 0.5 | 177 | 0.64 |
| Total water mass | 11.40 ± 1.18 | 11.98 ± 0.88 | -3.9 | 179 | $1.3 \times 10^{-4}$ |

**Table S4.** Linear effects of *Year* on standardized trait values for phenotypes measured in both years of the study. Bolded significant effects after Bonferroni correction for 33 models (p < 0.0015).

| | Phenotype | *n* | $\beta$ | SE | *p* | Adjusted $R^2$ |
| --- | --- | --- | --- | --- | --- | --- |
| **Capture** | **RMR** | 99 | **0.52** | **0.09** | **$3.0 \times 10^{-8}$** | 0.27 |
|  | **M<sub>sum</sub>** | 86 | **0.57** | **0.09** | **$8.9 \times 10^{-9}$** | 0.32 |
|  | Tarsus Length | 106 | -0.17 | 0.10 | 0.08 | 0.02 |
|  | M<sub>b</sub> | 106 | 0.04 | 0.10 | 0.69 | -0.01 |
|  | M<sub>lean</sub> | 98 | 0.03 | 0.10 | 0.80 | -0.01 |
|  | M<sub>fat</sub> | 98 | -0.10 | 0.10 | 0.24 | 0.00 |
|  | M<sub>fH2O</sub> | 98 | 0.02 | 0.10 | 0.82 | -0.01 |
| **Pre-acclimation** | **RMR** | 106 | **0.54** | **0.08** | **$1.8 \times 10^{-9}$** | 0.29 |
|  | M<sub>sum</sub> | 102 | -0.11 | 0.10 | 0.27 | 0.00 |
|  | **M<sub>b</sub>** | 106 | **0.42** | **0.09** | **$6.4 \times 10^{-6}$** | 0.17 |
|  | **M<sub>lean</sub>** | 105 | **-0.59** | **0.08** | **$2.1 \times 10^{-11}$** | 0.35 |
|  | **M<sub>fat</sub>** | 105 | **0.75** | **0.06** | **$< 2 \times 10^{-16}$** | 0.56 |
|  | M<sub>fH2O</sub> | 105 | -0.09 | 0.10 | 0.37 | 0.00 |
|  | **Hemoglobin** | 106 | **0.41** | **0.09** | **$9.4 \times 10^{-6}$** | 0.16 |
|  | Hematocrit | 100 | 0.16 | 0.10 | 0.10 | 0.02 |
| **Post-acclimation** | RMR | 105 | 0.11 | 0.10 | 0.25 | 0.00 |
|  | M<sub>sum</sub> | 105 | 0.15 | 0.10 | 0.13 | 0.01 |
|  | M<sub>b</sub> | 106 | -0.07 | 0.10 | 0.50 | -0.01 |
|  | M<sub>lean</sub> | 106 | -0.20 | 0.10 | 0.04 | 0.03 |
|  | M<sub>fat</sub> | 106 | 0.03 | 0.10 | 0.77 | -0.01 |
|  | M<sub>fH2O</sub> | 106 | 0.14 | 0.10 | 0.14 | 0.01 |
|  | Hemoglobin | 106 | -0.10 | 0.10 | 0.31 | 0.00 |
|  | Hematocrit | 105 | -0.25 | 0.09 | 0.01 | 0.05 |
|  | Erythrocyte count | 58 | -0.08 | 0.17 | 0.63 | -0.01 |
|  | **Heart mass** | 106 | **0.31** | **0.09** | **$1.1 \times 10^{-3}$** | 0.09 |
|  | Lung mass | 106 | 0.13 | 0.10 | 0.17 | 0.01 |
|  | Liver mass | 106 | 0.24 | 0.09 | 0.01 | 0.05 |
|  | Kidney mass | 104 | 0.16 | 0.10 | 0.10 | 0.02 |
|  | CPT | 95 | 0.18 | 0.10 | 0.08 | 0.02 |
|  | HOAD | 95 | 0.08 | 0.10 | 0.44 | 0.00 |
|  | CS | 95 | -0.03 | 0.10 | 0.76 | -0.01 |
|  | Plasma TRIG | 61 | 0.23 | 0.10 | 0.03 | 0.06 |
|  | Plasma glycerol | 64 | -0.11 | 0.13 | 0.39 | 0.00 |

**Table S5.** Pearson’s correlation coefficients for all pairwise trait associations at the capture.

| **Variable 1** | **Variable 2** | ***r*** | ***p*** |
| --- | --- | --- | --- |
| M<sub>sum</sub> | RMR | 0.38 | 4.1 x 10<sup>-4</sup> |
| M<sub>sum</sub> | Tarsus | 0.12 | 0.25 |
| RMR | Tarsus | 0.09 | 0.40 |
| M<sub>sum</sub> | Mass | 0.36 | 6.3 x 10<sup>-4</sup> |
| RMR | Mass | 0.54 | 1.3 x 10<sup>-8</sup> |
| Tarsus | Mass | 0.49 | 1.1 x 10<sup>-7</sup> |
| M<sub>sum</sub> | Lean | 0.28 | 0.01 |
| RMR | Lean | 0.44 | 1.1 x 10<sup>-5</sup> |
| Tarsus | Lean | 0.52 | 3.1 x 10<sup>-8</sup> |
| Mass | Lean | 0.96 | 8.1 x 10<sup>-57</sup> |
| M<sub>sum</sub> | Fat | 0.05 | 0.66 |
| RMR | Fat | 0.15 | 0.16 |
| Tarsus | Fat | 0.07 | 0.51 |
| Mass | Fat | 0.29 | 3.5 x 10<sup>-3</sup> |
| Lean | Fat | 0.24 | 0.02 |

**Table S6.** Pearson’s correlation coefficients for all pairwise trait associations at the end of the adjustment period (pre-acclimation).

| Variable 1 | Variable 2 | *r* | *p* |
| --- | --- | --- | --- |
| M<sub>sum</sub> | RMR | 0.41 | 2.0 x 10<sup>-5</sup> |
| M<sub>sum</sub> | Tarsus | -0.17 | 0.08 |
| RMR | Tarsus | -0.01 | 0.92 |
| M<sub>sum</sub> | Mass | 0.26 | 0.01 |
| RMR | Mass | 0.46 | 9.5 x 10<sup>-7</sup> |
| Tarsus | Mass | 0.14 | 0.15 |
| M<sub>sum</sub> | Lean | -0.31 | 1.4 x 10<sup>-3</sup> |
| RMR | Lean | -0.23 | 0.02 |
| Tarsus | Lean | 0.36 | 1.5 x 10<sup>-4</sup> |
| Mass | Lean | -0.04 | 0.70 |
| M<sub>sum</sub> | Fat | 0.45 | 2.1 x 10<sup>-6</sup> |
| RMR | Fat | 0.52 | 1.8 x 10<sup>-8</sup> |
| Tarsus | Fat | -0.12 | 0.22 |
| Mass | Fat | 0.83 | 2.0 x 10<sup>-27</sup> |
| Lean | Fat | -0.52 | 1.1 x 10<sup>-8</sup> |
| M<sub>sum</sub> | Hemoglobin | 0.27 | 6.9 x 10<sup>-3</sup> |
| RMR | Hemoglobin | 0.32 | 9.8 x 10<sup>-4</sup> |
| Tarsus | Hemoglobin | 0.02 | 0.86 |
| Mass | Hemoglobin | 0.33 | 5.2 x 10<sup>-4</sup> |
| Lean | Hemoglobin | -0.23 | 0.02 |
| Fat | Hemoglobin | 0.40 | 1.92 x 10<sup>-5</sup> |
| M<sub>sum</sub> | Hematocrit | 0.03 | 0.76 |
| RMR | Hematocrit | 0.11 | 0.28 |
| Tarsus | Hematocrit | 0.13 | 0.19 |
| Mass | Hematocrit | 0.25 | 0.01 |
| Lean | Hematocrit | -0.14 | 0.16 |
| Fat | Hematocrit | 0.26 | 0.01 |
| Hemoglobin | Hematocrit | 0.62 | 7.4 x 10<sup>-12</sup> |

**Table S7.**
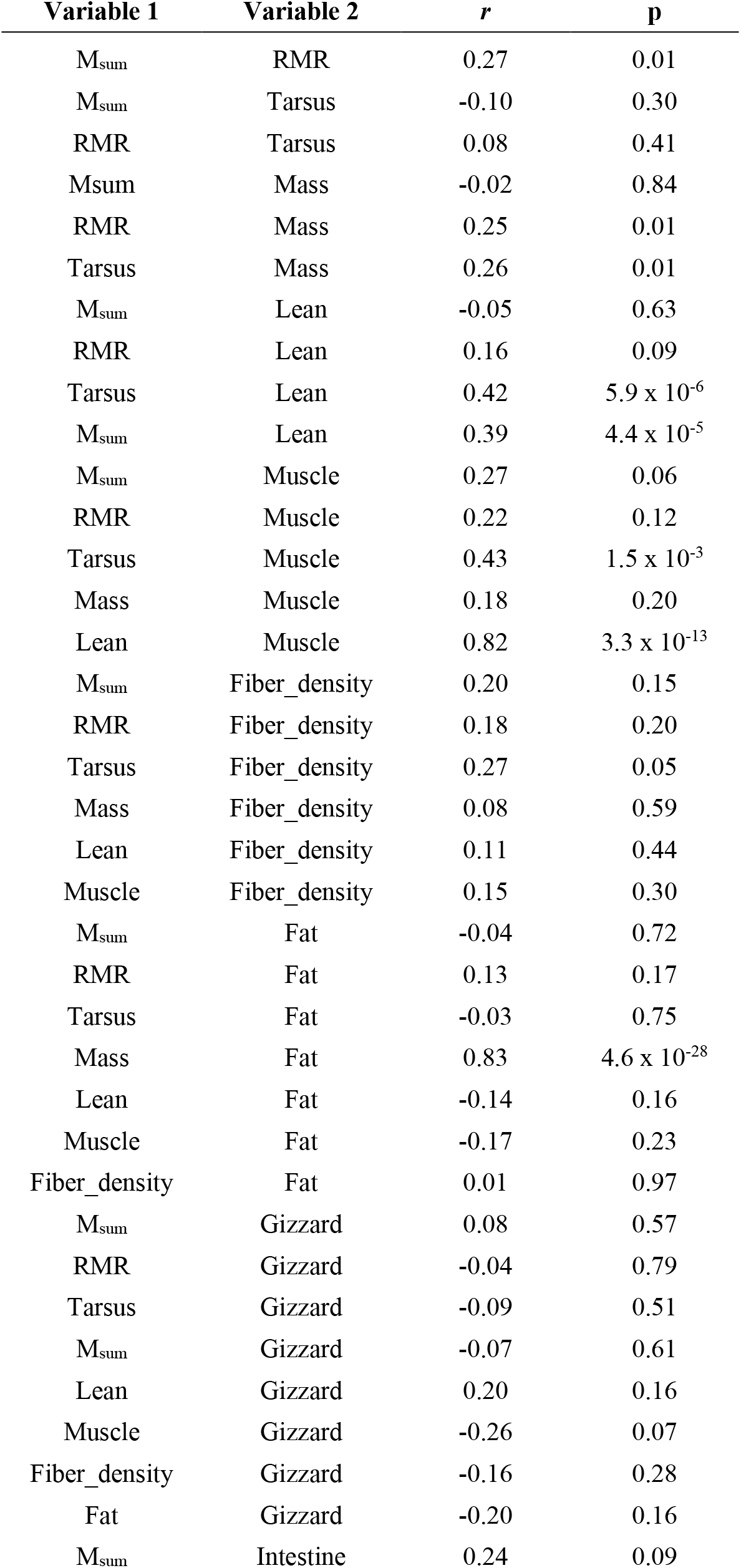
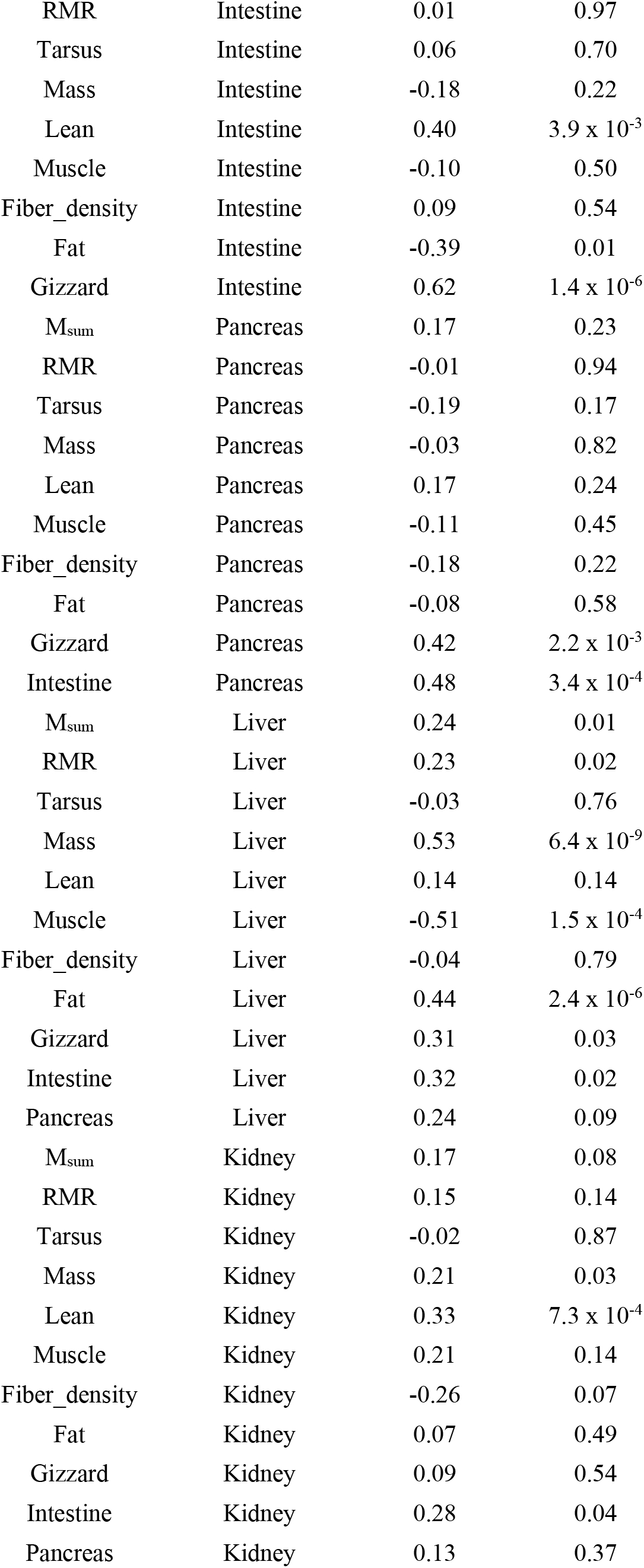
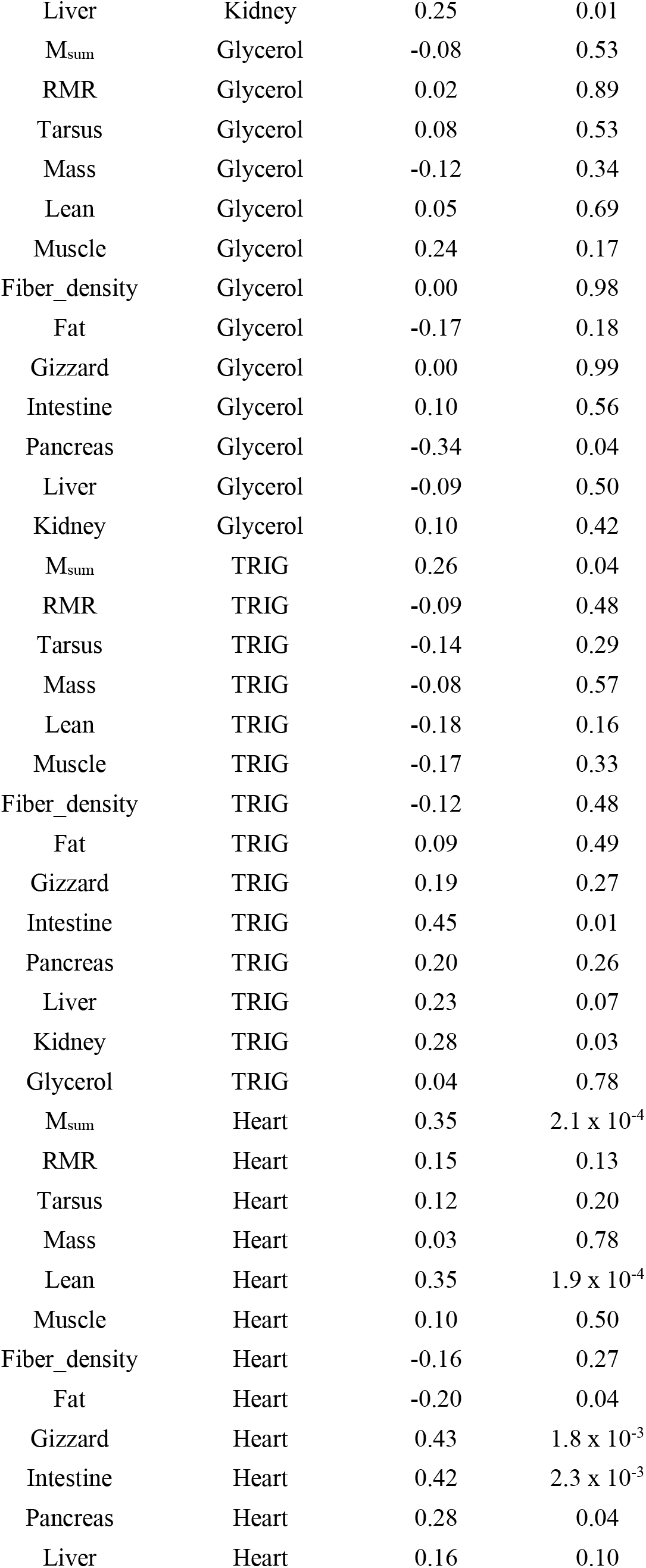
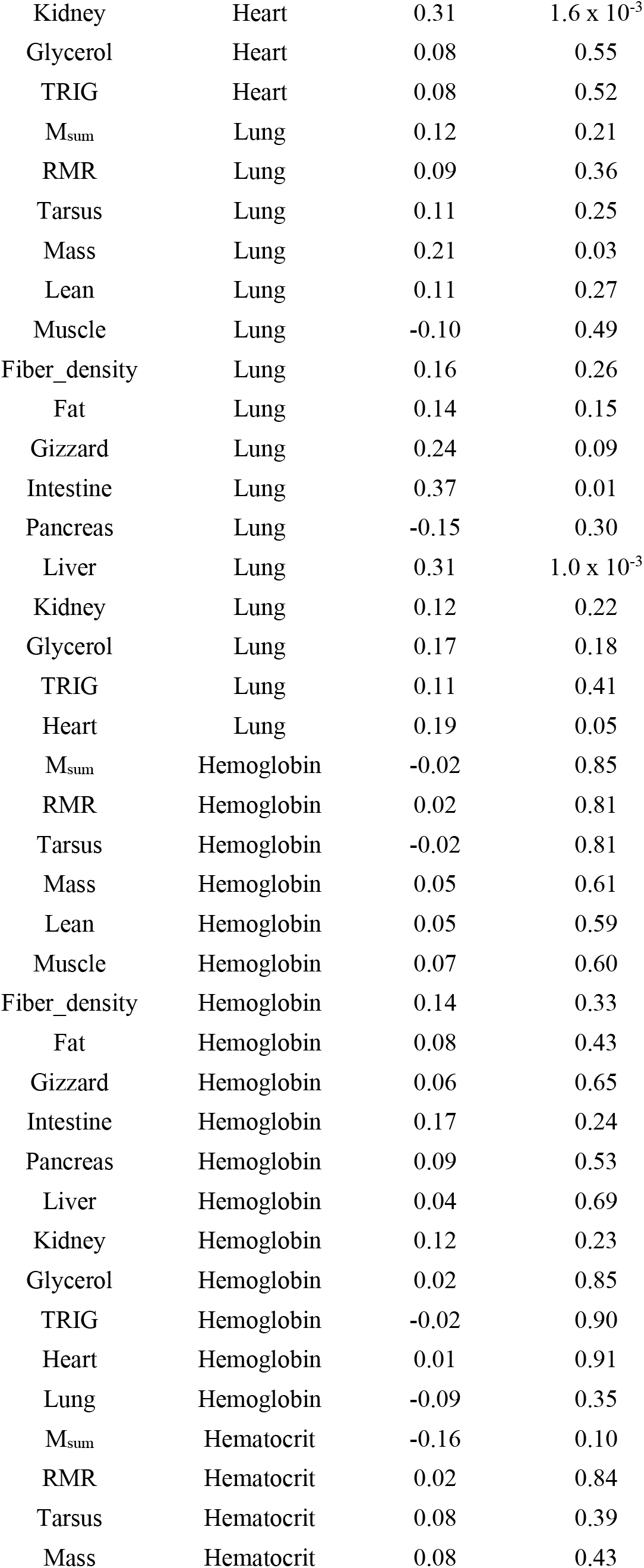
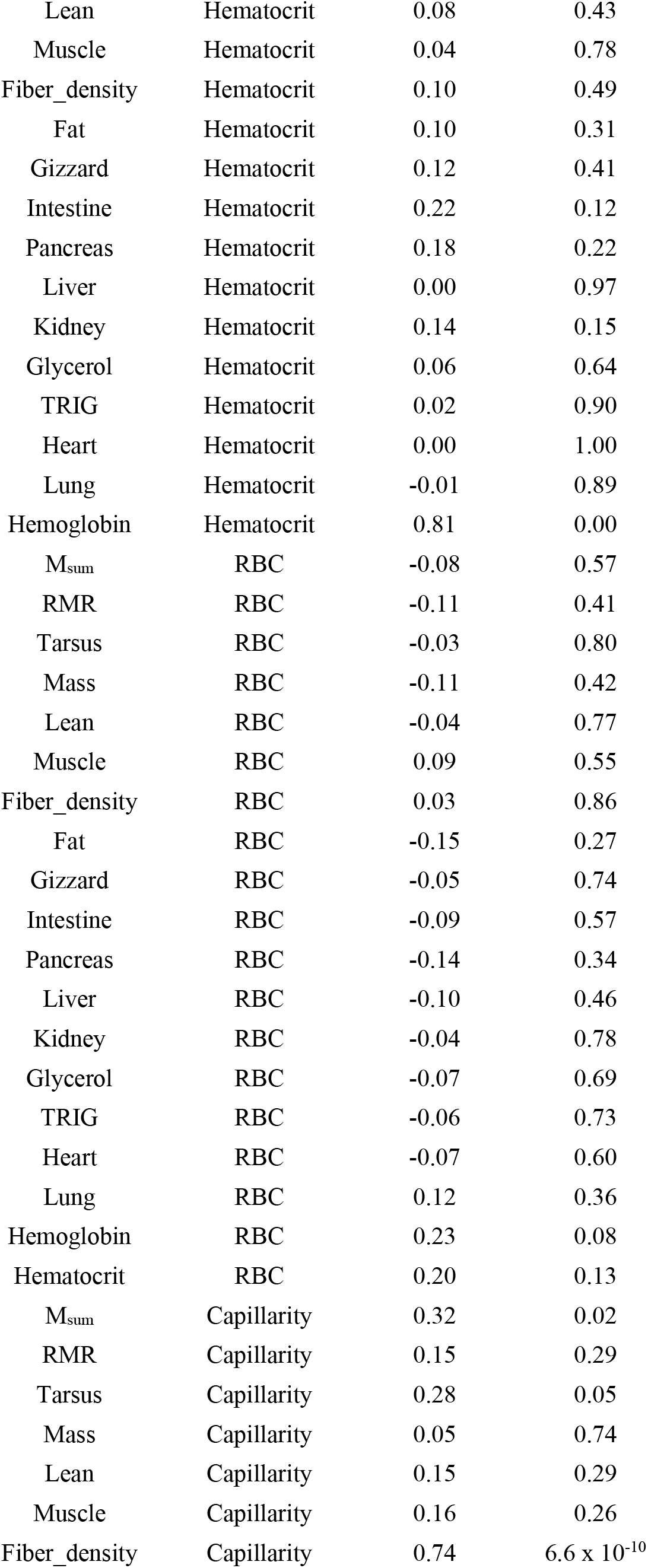
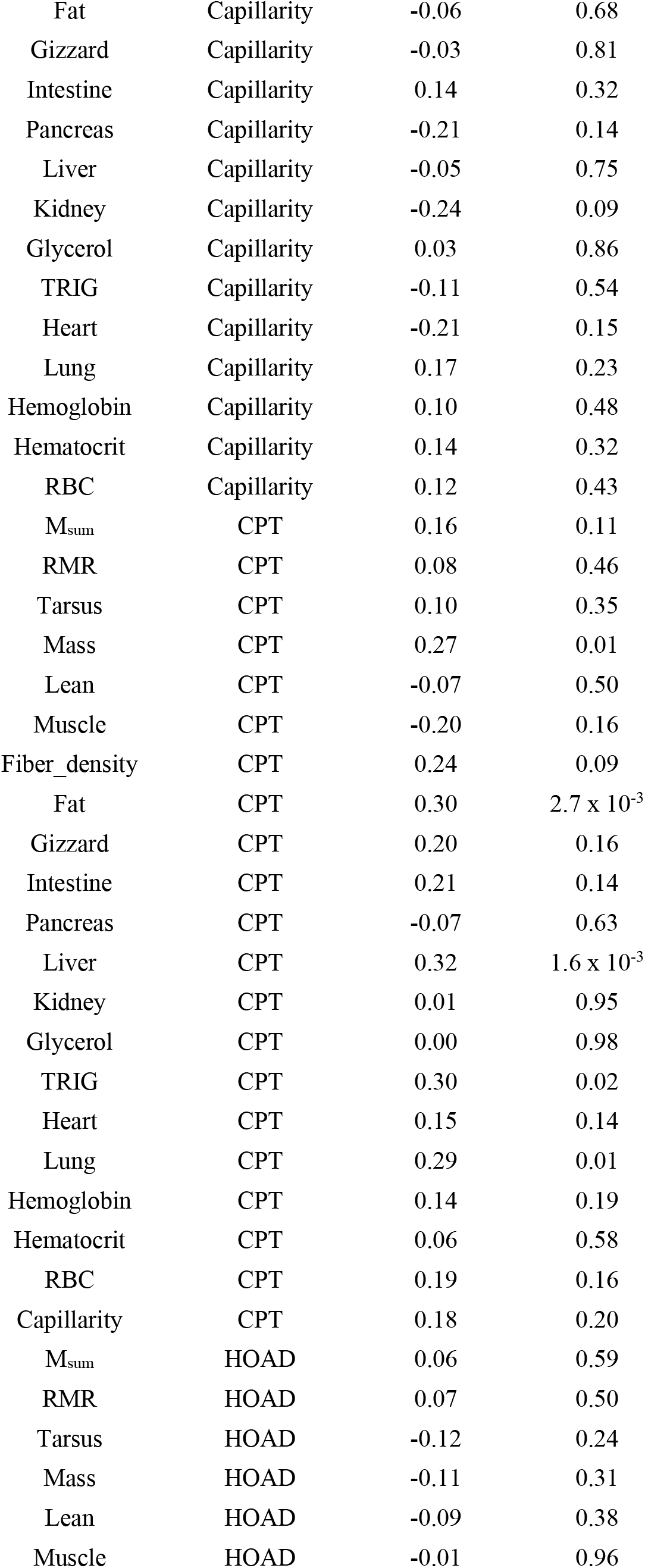
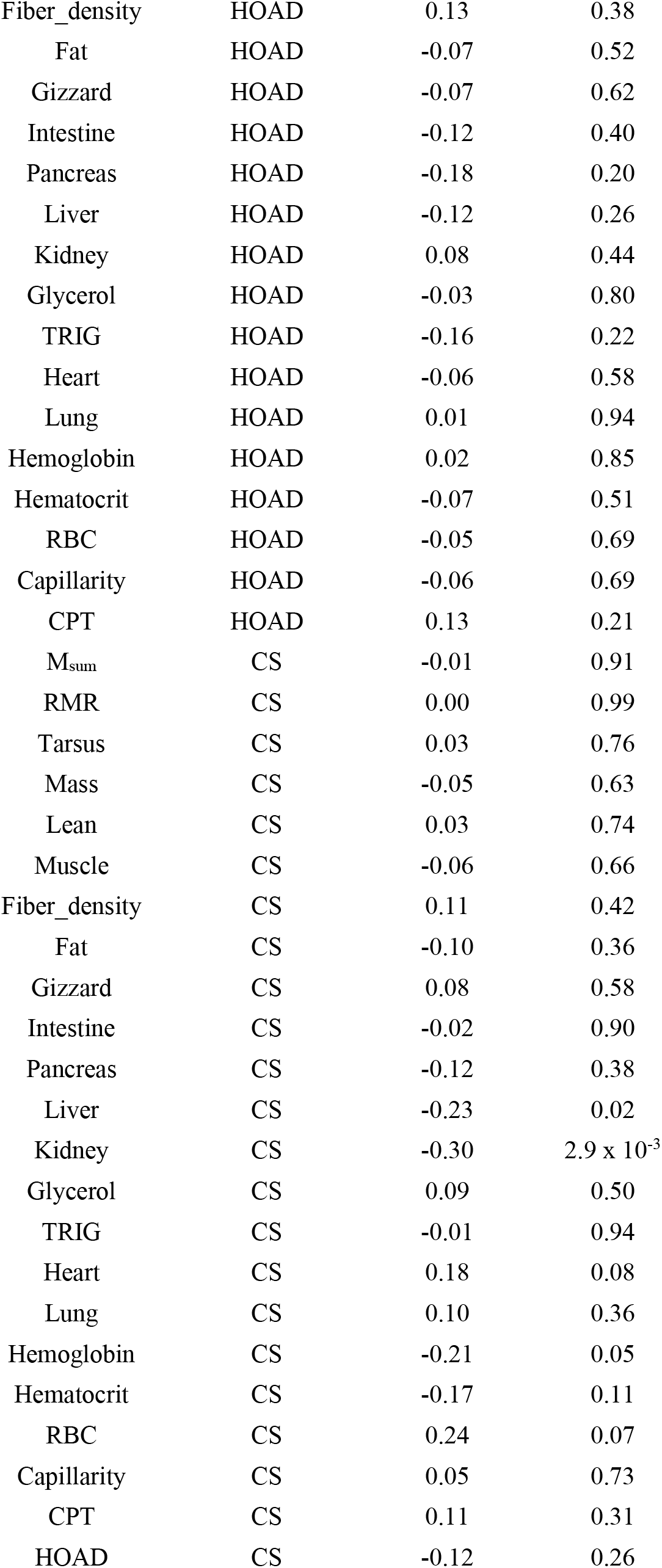
Pearson’s correlation coefficients for all pairwise trait associations post-acclimation.

| Variable 1 | Variable 2 | *r* | *p* |
| --- | --- | --- | --- |
| M<sub>sum</sub> | RMR | 0.27 | 0.01 |
| M<sub>sum</sub> | Tarsus | -0.10 | 0.30 |
| RMR | Tarsus | 0.08 | 0.41 |
| M<sub>sum</sub> | Mass | -0.02 | 0.84 |
| RMR | Mass | 0.25 | 0.01 |
| Tarsus | Mass | 0.26 | 0.01 |
| M<sub>sum</sub> | Lean | -0.05 | 0.63 |
| RMR | Lean | 0.16 | 0.09 |
| Tarsus | Lean | 0.42 | 5.9 x 10<sup>-6</sup> |
| M<sub>sum</sub> | Lean | 0.39 | 4.4 x 10<sup>-5</sup> |
| M<sub>sum</sub> | Muscle | 0.27 | 0.06 |
| RMR | Muscle | 0.22 | 0.12 |
| Tarsus | Muscle | 0.43 | 1.5 x 10<sup>-3</sup> |
| Mass | Muscle | 0.18 | 0.20 |
| Lean | Muscle | 0.82 | 3.3 x 10<sup>-13</sup> |
| M<sub>sum</sub> | Fiber_density | 0.20 | 0.15 |
| RMR | Fiber_density | 0.18 | 0.20 |
| Tarsus | Fiber_density | 0.27 | 0.05 |
| Mass | Fiber_density | 0.08 | 0.59 |
| Lean | Fiber_density | 0.11 | 0.44 |
| Muscle | Fiber_density | 0.15 | 0.30 |
| M<sub>sum</sub> | Fat | -0.04 | 0.72 |
| RMR | Fat | 0.13 | 0.17 |
| Tarsus | Fat | -0.03 | 0.75 |
| Mass | Fat | 0.83 | 4.6 x 10<sup>-28</sup> |
| Lean | Fat | -0.14 | 0.16 |
| Muscle | Fat | -0.17 | 0.23 |
| Fiber_density | Fat | 0.01 | 0.97 |
| M<sub>sum</sub> | Gizzard | 0.08 | 0.57 |
| RMR | Gizzard | -0.04 | 0.79 |
| Tarsus | Gizzard | -0.09 | 0.51 |
| M<sub>sum</sub> | Gizzard | -0.07 | 0.61 |
| Lean | Gizzard | 0.20 | 0.16 |
| Muscle | Gizzard | -0.26 | 0.07 |
| Fiber_density | Gizzard | -0.16 | 0.28 |
| Fat | Gizzard | -0.20 | 0.16 |
| M<sub>sum</sub> | Intestine | 0.24 | 0.09 |
| RMR | Intestine | 0.01 | 0.97 |
| Tarsus | Intestine | 0.06 | 0.70 |
| Mass | Intestine | -0.18 | 0.22 |
| Lean | Intestine | 0.40 | $3.9 \times 10^{-3}$ |
| Muscle | Intestine | -0.10 | 0.50 |
| Fiber_density | Intestine | 0.09 | 0.54 |
| Fat | Intestine | -0.39 | 0.01 |
| Gizzard | Intestine | 0.62 | $1.4 \times 10^{-6}$ |
| M<sub>sum</sub> | Pancreas | 0.17 | 0.23 |
| RMR | Pancreas | -0.01 | 0.94 |
| Tarsus | Pancreas | -0.19 | 0.17 |
| Mass | Pancreas | -0.03 | 0.82 |
| Lean | Pancreas | 0.17 | 0.24 |
| Muscle | Pancreas | -0.11 | 0.45 |
| Fiber_density | Pancreas | -0.18 | 0.22 |
| Fat | Pancreas | -0.08 | 0.58 |
| Gizzard | Pancreas | 0.42 | $2.2 \times 10^{-3}$ |
| Intestine | Pancreas | 0.48 | $3.4 \times 10^{-4}$ |
| M<sub>sum</sub> | Liver | 0.24 | 0.01 |
| RMR | Liver | 0.23 | 0.02 |
| Tarsus | Liver | -0.03 | 0.76 |
| Mass | Liver | 0.53 | $6.4 \times 10^{-9}$ |
| Lean | Liver | 0.14 | 0.14 |
| Muscle | Liver | -0.51 | $1.5 \times 10^{-4}$ |
| Fiber_density | Liver | -0.04 | 0.79 |
| Fat | Liver | 0.44 | $2.4 \times 10^{-6}$ |
| Gizzard | Liver | 0.31 | 0.03 |
| Intestine | Liver | 0.32 | 0.02 |
| Pancreas | Liver | 0.24 | 0.09 |
| M<sub>sum</sub> | Kidney | 0.17 | 0.08 |
| RMR | Kidney | 0.15 | 0.14 |
| Tarsus | Kidney | -0.02 | 0.87 |
| Mass | Kidney | 0.21 | 0.03 |
| Lean | Kidney | 0.33 | $7.3 \times 10^{-4}$ |
| Muscle | Kidney | 0.21 | 0.14 |
| Fiber_density | Kidney | -0.26 | 0.07 |
| Fat | Kidney | 0.07 | 0.49 |
| Gizzard | Kidney | 0.09 | 0.54 |
| Intestine | Kidney | 0.28 | 0.04 |
| Pancreas | Kidney | 0.13 | 0.37 |
| Liver | Kidney | 0.25 | 0.01 |
| M<sub>sum</sub> | Glycerol | -0.08 | 0.53 |
| RMR | Glycerol | 0.02 | 0.89 |
| Tarsus | Glycerol | 0.08 | 0.53 |
| Mass | Glycerol | -0.12 | 0.34 |
| Lean | Glycerol | 0.05 | 0.69 |
| Muscle | Glycerol | 0.24 | 0.17 |
| Fiber_density | Glycerol | 0.00 | 0.98 |
| Fat | Glycerol | -0.17 | 0.18 |
| Gizzard | Glycerol | 0.00 | 0.99 |
| Intestine | Glycerol | 0.10 | 0.56 |
| Pancreas | Glycerol | -0.34 | 0.04 |
| Liver | Glycerol | -0.09 | 0.50 |
| Kidney | Glycerol | 0.10 | 0.42 |
| M<sub>sum</sub> | TRIG | 0.26 | 0.04 |
| RMR | TRIG | -0.09 | 0.48 |
| Tarsus | TRIG | -0.14 | 0.29 |
| Mass | TRIG | -0.08 | 0.57 |
| Lean | TRIG | -0.18 | 0.16 |
| Muscle | TRIG | -0.17 | 0.33 |
| Fiber_density | TRIG | -0.12 | 0.48 |
| Fat | TRIG | 0.09 | 0.49 |
| Gizzard | TRIG | 0.19 | 0.27 |
| Intestine | TRIG | 0.45 | 0.01 |
| Pancreas | TRIG | 0.20 | 0.26 |
| Liver | TRIG | 0.23 | 0.07 |
| Kidney | TRIG | 0.28 | 0.03 |
| Glycerol | TRIG | 0.04 | 0.78 |
| M<sub>sum</sub> | Heart | 0.35 | 2.1 x 10<sup>-4</sup> |
| RMR | Heart | 0.15 | 0.13 |
| Tarsus | Heart | 0.12 | 0.20 |
| Mass | Heart | 0.03 | 0.78 |
| Lean | Heart | 0.35 | 1.9 x 10<sup>-4</sup> |
| Muscle | Heart | 0.10 | 0.50 |
| Fiber_density | Heart | -0.16 | 0.27 |
| Fat | Heart | -0.20 | 0.04 |
| Gizzard | Heart | 0.43 | 1.8 x 10<sup>-3</sup> |
| Intestine | Heart | 0.42 | 2.3 x 10<sup>-3</sup> |
| Pancreas | Heart | 0.28 | 0.04 |
| Liver | Heart | 0.16 | 0.10 |
| Kidney | Heart | 0.31 | $1.6 \times 10^{-3}$ |
| Glycerol | Heart | 0.08 | 0.55 |
| TRIG | Heart | 0.08 | 0.52 |
| M<sub>sum</sub> | Lung | 0.12 | 0.21 |
| RMR | Lung | 0.09 | 0.36 |
| Tarsus | Lung | 0.11 | 0.25 |
| Mass | Lung | 0.21 | 0.03 |
| Lean | Lung | 0.11 | 0.27 |
| Muscle | Lung | -0.10 | 0.49 |
| Fiber_density | Lung | 0.16 | 0.26 |
| Fat | Lung | 0.14 | 0.15 |
| Gizzard | Lung | 0.24 | 0.09 |
| Intestine | Lung | 0.37 | 0.01 |
| Pancreas | Lung | -0.15 | 0.30 |
| Liver | Lung | 0.31 | $1.0 \times 10^{-3}$ |
| Kidney | Lung | 0.12 | 0.22 |
| Glycerol | Lung | 0.17 | 0.18 |
| TRIG | Lung | 0.11 | 0.41 |
| Heart | Lung | 0.19 | 0.05 |
| M<sub>sum</sub> | Hemoglobin | -0.02 | 0.85 |
| RMR | Hemoglobin | 0.02 | 0.81 |
| Tarsus | Hemoglobin | -0.02 | 0.81 |
| Mass | Hemoglobin | 0.05 | 0.61 |
| Lean | Hemoglobin | 0.05 | 0.59 |
| Muscle | Hemoglobin | 0.07 | 0.60 |
| Fiber_density | Hemoglobin | 0.14 | 0.33 |
| Fat | Hemoglobin | 0.08 | 0.43 |
| Gizzard | Hemoglobin | 0.06 | 0.65 |
| Intestine | Hemoglobin | 0.17 | 0.24 |
| Pancreas | Hemoglobin | 0.09 | 0.53 |
| Liver | Hemoglobin | 0.04 | 0.69 |
| Kidney | Hemoglobin | 0.12 | 0.23 |
| Glycerol | Hemoglobin | 0.02 | 0.85 |
| TRIG | Hemoglobin | -0.02 | 0.90 |
| Heart | Hemoglobin | 0.01 | 0.91 |
| Lung | Hemoglobin | -0.09 | 0.35 |
| M<sub>sum</sub> | Hematocrit | -0.16 | 0.10 |
| RMR | Hematocrit | 0.02 | 0.84 |
| Tarsus | Hematocrit | 0.08 | 0.39 |
| Mass | Hematocrit | 0.08 | 0.43 |
| Lean | Hematocrit | 0.08 | 0.43 |
| Muscle | Hematocrit | 0.04 | 0.78 |
| Fiber_density | Hematocrit | 0.10 | 0.49 |
| Fat | Hematocrit | 0.10 | 0.31 |
| Gizzard | Hematocrit | 0.12 | 0.41 |
| Intestine | Hematocrit | 0.22 | 0.12 |
| Pancreas | Hematocrit | 0.18 | 0.22 |
| Liver | Hematocrit | 0.00 | 0.97 |
| Kidney | Hematocrit | 0.14 | 0.15 |
| Glycerol | Hematocrit | 0.06 | 0.64 |
| TRIG | Hematocrit | 0.02 | 0.90 |
| Heart | Hematocrit | 0.00 | 1.00 |
| Lung | Hematocrit | -0.01 | 0.89 |
| Hemoglobin | Hematocrit | 0.81 | 0.00 |
| M<sub>sum</sub> | RBC | -0.08 | 0.57 |
| RMR | RBC | -0.11 | 0.41 |
| Tarsus | RBC | -0.03 | 0.80 |
| Mass | RBC | -0.11 | 0.42 |
| Lean | RBC | -0.04 | 0.77 |
| Muscle | RBC | 0.09 | 0.55 |
| Fiber_density | RBC | 0.03 | 0.86 |
| Fat | RBC | -0.15 | 0.27 |
| Gizzard | RBC | -0.05 | 0.74 |
| Intestine | RBC | -0.09 | 0.57 |
| Pancreas | RBC | -0.14 | 0.34 |
| Liver | RBC | -0.10 | 0.46 |
| Kidney | RBC | -0.04 | 0.78 |
| Glycerol | RBC | -0.07 | 0.69 |
| TRIG | RBC | -0.06 | 0.73 |
| Heart | RBC | -0.07 | 0.60 |
| Lung | RBC | 0.12 | 0.36 |
| Hemoglobin | RBC | 0.23 | 0.08 |
| Hematocrit | RBC | 0.20 | 0.13 |
| M<sub>sum</sub> | Capillarity | 0.32 | 0.02 |
| RMR | Capillarity | 0.15 | 0.29 |
| Tarsus | Capillarity | 0.28 | 0.05 |
| Mass | Capillarity | 0.05 | 0.74 |
| Lean | Capillarity | 0.15 | 0.29 |
| Muscle | Capillarity | 0.16 | 0.26 |
| Fiber_density | Capillarity | 0.74 | 6.6 x 10<sup>-10</sup> |
| Fat | Capillarity | -0.06 | 0.68 |
| Gizzard | Capillarity | -0.03 | 0.81 |
| Intestine | Capillarity | 0.14 | 0.32 |
| Pancreas | Capillarity | -0.21 | 0.14 |
| Liver | Capillarity | -0.05 | 0.75 |
| Kidney | Capillarity | -0.24 | 0.09 |
| Glycerol | Capillarity | 0.03 | 0.86 |
| TRIG | Capillarity | -0.11 | 0.54 |
| Heart | Capillarity | -0.21 | 0.15 |
| Lung | Capillarity | 0.17 | 0.23 |
| Hemoglobin | Capillarity | 0.10 | 0.48 |
| Hematocrit | Capillarity | 0.14 | 0.32 |
| RBC | Capillarity | 0.12 | 0.43 |
| M<sub>sum</sub> | CPT | 0.16 | 0.11 |
| RMR | CPT | 0.08 | 0.46 |
| Tarsus | CPT | 0.10 | 0.35 |
| Mass | CPT | 0.27 | 0.01 |
| Lean | CPT | -0.07 | 0.50 |
| Muscle | CPT | -0.20 | 0.16 |
| Fiber_density | CPT | 0.24 | 0.09 |
| Fat | CPT | 0.30 | 2.7 x 10<sup>-3</sup> |
| Gizzard | CPT | 0.20 | 0.16 |
| Intestine | CPT | 0.21 | 0.14 |
| Pancreas | CPT | -0.07 | 0.63 |
| Liver | CPT | 0.32 | 1.6 x 10<sup>-3</sup> |
| Kidney | CPT | 0.01 | 0.95 |
| Glycerol | CPT | 0.00 | 0.98 |
| TRIG | CPT | 0.30 | 0.02 |
| Heart | CPT | 0.15 | 0.14 |
| Lung | CPT | 0.29 | 0.01 |
| Hemoglobin | CPT | 0.14 | 0.19 |
| Hematocrit | CPT | 0.06 | 0.58 |
| RBC | CPT | 0.19 | 0.16 |
| Capillarity | CPT | 0.18 | 0.20 |
| M<sub>sum</sub> | HOAD | 0.06 | 0.59 |
| RMR | HOAD | 0.07 | 0.50 |
| Tarsus | HOAD | -0.12 | 0.24 |
| Mass | HOAD | -0.11 | 0.31 |
| Lean | HOAD | -0.09 | 0.38 |
| Muscle | HOAD | -0.01 | 0.96 |
| Fiber_density | HOAD | 0.13 | 0.38 |
| Fat | HOAD | -0.07 | 0.52 |
| Gizzard | HOAD | -0.07 | 0.62 |
| Intestine | HOAD | -0.12 | 0.40 |
| Pancreas | HOAD | -0.18 | 0.20 |
| Liver | HOAD | -0.12 | 0.26 |
| Kidney | HOAD | 0.08 | 0.44 |
| Glycerol | HOAD | -0.03 | 0.80 |
| TRIG | HOAD | -0.16 | 0.22 |
| Heart | HOAD | -0.06 | 0.58 |
| Lung | HOAD | 0.01 | 0.94 |
| Hemoglobin | HOAD | 0.02 | 0.85 |
| Hematocrit | HOAD | -0.07 | 0.51 |
| RBC | HOAD | -0.05 | 0.69 |
| Capillarity | HOAD | -0.06 | 0.69 |
| CPT | HOAD | 0.13 | 0.21 |
| M<sub>sum</sub> | CS | -0.01 | 0.91 |
| RMR | CS | 0.00 | 0.99 |
| Tarsus | CS | 0.03 | 0.76 |
| Mass | CS | -0.05 | 0.63 |
| Lean | CS | 0.03 | 0.74 |
| Muscle | CS | -0.06 | 0.66 |
| Fiber_density | CS | 0.11 | 0.42 |
| Fat | CS | -0.10 | 0.36 |
| Gizzard | CS | 0.08 | 0.58 |
| Intestine | CS | -0.02 | 0.90 |
| Pancreas | CS | -0.12 | 0.38 |
| Liver | CS | -0.23 | 0.02 |
| Kidney | CS | -0.30 | $2.9 \times 10^{-3}$ |
| Glycerol | CS | 0.09 | 0.50 |
| TRIG | CS | -0.01 | 0.94 |
| Heart | CS | 0.18 | 0.08 |
| Lung | CS | 0.10 | 0.36 |
| Hemoglobin | CS | -0.21 | 0.05 |
| Hematocrit | CS | -0.17 | 0.11 |
| RBC | CS | 0.24 | 0.07 |
| Capillarity | CS | 0.05 | 0.73 |
| CPT | CS | 0.11 | 0.31 |
| HOAD | CS | -0.12 | 0.26 |

